# Polymorphic Inverted Repeats near coding genes impact chromatin topology and phenotypic traits in *Arabidopsis thaliana*

**DOI:** 10.1101/2022.07.05.498814

**Authors:** Agustín L. Arce, Regina Mencia, Damian A. Cambiagno, Patricia L. Lang, Chang Liu, Hernán A. Burbano, Detlef Weigel, Pablo A. Manavella

**Author notes:** These authors contributed equally (Agustín L. Arce, Regina Mencia).

## Abstract

Transposons are mobile elements that are commonly silenced to protect eukaryotic genome integrity. In plants, transposable elements (TEs) can be activated during stress conditions and subsequently insert into gene-rich regions. TE-derived inverted repeats (IRs) are commonly found near plant genes, where they affect host gene expression with potentially positive effects on adaptation. However, the molecular mechanisms by which these IRs control gene expression is unclear in most cases. Here, we identify in the *Arabidopsis thaliana* genome hundreds of IRs located near genes that are transcribed by RNA Polymerase II, resulting in the production of 24-nt small RNAs that trigger methylation of the IRs. The expression of these IRs is associated with drastic changes in the local 3D chromatin organization, which alter the expression pattern of the hosting genes. Notably, the presence and structure of many IRs differ between *A. thaliana* accessions. Capture-C sequencing experiments revealed that such variation changes short-range chromatin interactions, which translates into changes in gene expression patterns. CRISPR/Cas9-mediated disruption of two of such IRs leads to a switch in genome topology and gene expression, with phenotypic consequences. Our data demonstrate that the insertion of an IR near a gene provides an anchor point for chromatin interactions that can profoundly impact the activity of neighboring loci. This turns IRs into powerful evolutionary agents that can contribute to rapid adaptation.

## Introduction

Transposable elements (TEs) are widely distributed among eukaryotic genomes. In a process known as transposition, TEs move within the genome to different locations, usually copying themself as they ‘jump’ (Dubin et al., 2018). Plant genomes are particularly rich in TEs and repetitive elements, which, for example, account for 85% of the maize genome (Schnable et al., 2009). In plants, TEs are commonly silenced through DNA methylation in a process known as RNA-directed DNA methylation (RdDM), which maintains genome integrity (Matzke et al., 2015). To trigger RdDM, short RNA Polymerase IV (RNAPIV)-dependent TE transcripts are converted into double-stranded RNA (dsRNA) by the RNA-dependent RNA polymerase 2 (RDR2) and then to 24 nt small interfering RNAs (siRNAs) by DICER-Like 3 (DCL3) (Matzke *et al*., 2015; Zhou and Law, 2015). ARGONAUTE 4 (AGO4)-loaded siRNAs then direct de novo methylation of the TE loci by recognizing nascent RNAPV transcripts there. Such methylation ultimately leads to nucleosome condensation and permanent silencing of the TE. Still, massive bursts of TE amplification have occurred in plant genomes in addition to the rarer, but continued movement of individual elements (Lu et al., 2012; Maumus and Quesneville, 2014; SanMiguel et al., 1998). Stress can trigger the activation of TEs and fuel transposition (Baduel et al., 2021; Tittel-Elmer et al., 2010). TEs are thus significant contributors to genetic variation in plant genomes (Baduel *et al*., 2021; Collier and Largaespada, 2007; Deragon et al., 2008) and have been postulated as drivers of genome evolution and expansion, as well as developmental plasticity and adaptation (Dubin *et al*., 2018; Lisch, 2013).

There are two main classes of TEs: DNA transposons and retrotransposons. The most abundant DNA transposons are miniature inverted-repeat TEs (MITEs), while the most abundant retrotransposons are long terminal repeat retrotransposons (LTRs) (Dubin *et al*., 2018). MITEs exhibit characteristic terminal inverted repeats (TIRs) and small direct repeats (target site duplications, TSDs), but lack transposase sequences, making them non-autonomous elements (Fattash et al., 2013; Yang et al., 2009). Many TE-derived inverted repeated (IR) insertions may not be classified as MITE as they lack the above-mentioned components of MITEs, either due to deletions after insertion, or because they were generated through a different process. MITEs are commonly situated near coding genes: for example, almost 60% of rice genes can be associated with a MITE (Lu *et al*., 2012), with the MITEs often changing the expression of neighboring genes (Lu *et al*., 2012; Underwood et al., 2022; Wu et al., 2022; Xu et al., 2020; Zhang et al., 2016). Based on this, MITEs have been proposed to play important roles in genome evolution and gene expression (Lu *et al*., 2012). One key feature of MITEs is that their transcripts can fold into hairpin-shaped dsRNAs due to the extensive sequence complementarity between IR arms. These dsRNA secondary structures are recognized and processed by DCL3 to produce 24-nt siRNAs that trigger DNA methylation without the need for RNAPIV/RDR2 activity (Ariel and Manavella, 2021; Crescente et al., 2022; Cuerda-Gil and Slotkin, 2016; Gagliardi et al., 2019; Sasaki et al., 2014). Thus, transcripts of these MITEs can be initiated from promoters of adjacent genes, triggering their RNA Polymerase II (RNAPII)-dependent DNA methylation. At the sunflower *HaWRKY6* locus, the RNAPII-mediated transcription of a MITE triggers the methylation of its coding region and causes the formation of alternative regulatory short-range chromatin loops in the locus that specifically change its expression (Gagliardi *et al*., 2019).

Three-dimensional chromatin organization has recently emerged as a critical feature determining genome functionality, fine-tuning gene expression and developmental responses in plants (Domb et al., 2022; Zhang and Wang, 2021). Short-range chromatin loops reflect interaction between relatively close regions of DNA, within a few kb, generally within a single locus or between adjacent loci (Gagliardi and Manavella, 2020). Different from canonical regulation by mC methylation in the linear DNA sequence, commonly associated with gene repression, local three-dimensional chromatin organization can induce a plethora of regulatory mechanisms, including transcriptional activation/repression, transcription directionality, alternative splicing, the usage of cryptic termination sites, impaired or enhanced RNAPII elongation, DNA replication and repair (Gagliardi and Manavella, 2020; Grzechnik et al., 2014; Sotelo-Silveira et al., 2018).

In this study, we show that TE-derived IR elements located near genes in the *Arabidopsis thaliana* genome can cause rearrangements of the chromatin topology, promoting the formation of short-range chromatin loops. These chromatin interactions, which depend on the production of IR-derived siRNAs and *de novo* DNA methylation, often translate into changes in gene expression. The presence of an IR and its associated chromatin loop near a gene does not cause a uniform regulatory effect, and can either enhance or repress expression depending on the locus and which regions within the locus are part of the loop. Almost one third of the identified gene-associated IRs are not conserved among a set of 216 *A. thaliana* natural accessions. Our data show that polymorphisms in IRs near genes can be coupled with a change in the chromatin topology of the region. These accession-related changes in chromatin landscapes controlled by IRs can be linked to alteration in traits commonly associated with adaptation, such as flowering. In proof-of-concept experiments, we used CRISPR/Cas9 genome editing to mimic the situation in natural accessions lacking specific IRs, and demonstrated that the IRs help to shape chromatin topology, which in turn can control molecular and organismal phenotypes. We found that IRs downstream of *PHYC* and *CRY1* cause the formation of repressive chromatin loops associated with well-defined developmental phenotypes of some natural accessions. Overall, our data demonstrate that TE-derived IRs can produce changes in chromatin topology, gene expression, and ultimately phenotypic changes through their capacity to trigger DNA methylation autonomously. Given the propensity of TEs to activate during stress responses and the tendency of IRs to locate near coding genes, our finding provides a scenario for these TEs to drive local adaptation and domestication by supporting rapid and sometimes drastic changes in 3D chromatin organization and gene activity after single-set mutational events.

## Results

### TE-derived IRs located near genes impact the local chromatin topology in Arabidopsis

A TE-derived IR element located ∼600 bp upstream of the *HaWRKY6* locus in sunflower serves as an anchor point for the formation of two short-range chromatin loops, and thereby promotes changes in local chromatin topology (Gagliardi *et al*., 2019). The ultimate outcome is the methylation of the locus, due to 24-nt siRNAs produced after RNAPII-dependent transcription of these IRs. In leaves, this leads to the formation of a repressive loop, while it promotes a second, larger, loop that enhances transcription in cotyledons (Gagliardi *et al*., 2019).

Because TE-derived IRs are frequently located near genes (Guo et al., 2017), we wondered whether the *HaWRKY6* case was just one example of a more general phenomenon in plants. To evaluate if the insertion of an IR near a gene changes local chromatin topology and gene expression, we first aimed to identify all Irs neighboring protein-coding genes in the *A. thaliana* Col-0 reference genome. Using *einverted* from the EMBOSS program suite (Rice et al., 2000), we found a total of 885 IRs in the *A. thaliana* genome, 634 of then near annotated protein coding genes (222 of which were located within 500 bp upstream or downstream of a protein coding gene, a further 163 between 500 bp and 1,000 bp from a gene, and 249 between 1,000 and 3,000 bp from a gene) (Fig. 1A). IRs were found within 500 bp of 260 unique genes, within 500 to 1,000 bp of 215 unique genes, and within 1,000 to 3,000 bp of 615 unique genes (Fig. 1A). These IRs have a broad genome-wide distribution, with many located in the gene-rich chromosome arms, and others in gene-poor/TE-rich pericentromeric regions (Fig. 1B). Analyzing the overlap with annotated transposable elements (TEs) revealed that 68% of the IRs within 3,000 bp of a gene were clearly of TE origin, with most (∼44%) from the MuDR superfamily and 18% from the Helitron superfamily (Fig. 1C). These percentages remain invariable despite of the distance from the IR to the hosting gene although a moderate enrichment is LTR/Gypsy TEs between IRs not associated with genes (Fig. 1C). Using the plaNETseq dataset of RNAPII associated nascent transcripts (Kindgren et al., 2020), we found that more than half of the identified gene-associated IRs are transcribed by RNAPII (Fig. 1D). Supporting the idea that these IRs are transcribed from promoters of nearby genes, the fraction of IRs transcribed by RNAPII increases the closer the IRs are to a gene (Fig. 1D). Conversely, IRs located far from annotated genes were less likely to give rise to RNAPII transcripts. Instead, their transcription likely depends on the canonical RNAPIV/RDR2 RdDM pathway (Fig. 1D).

**Figure 1.**
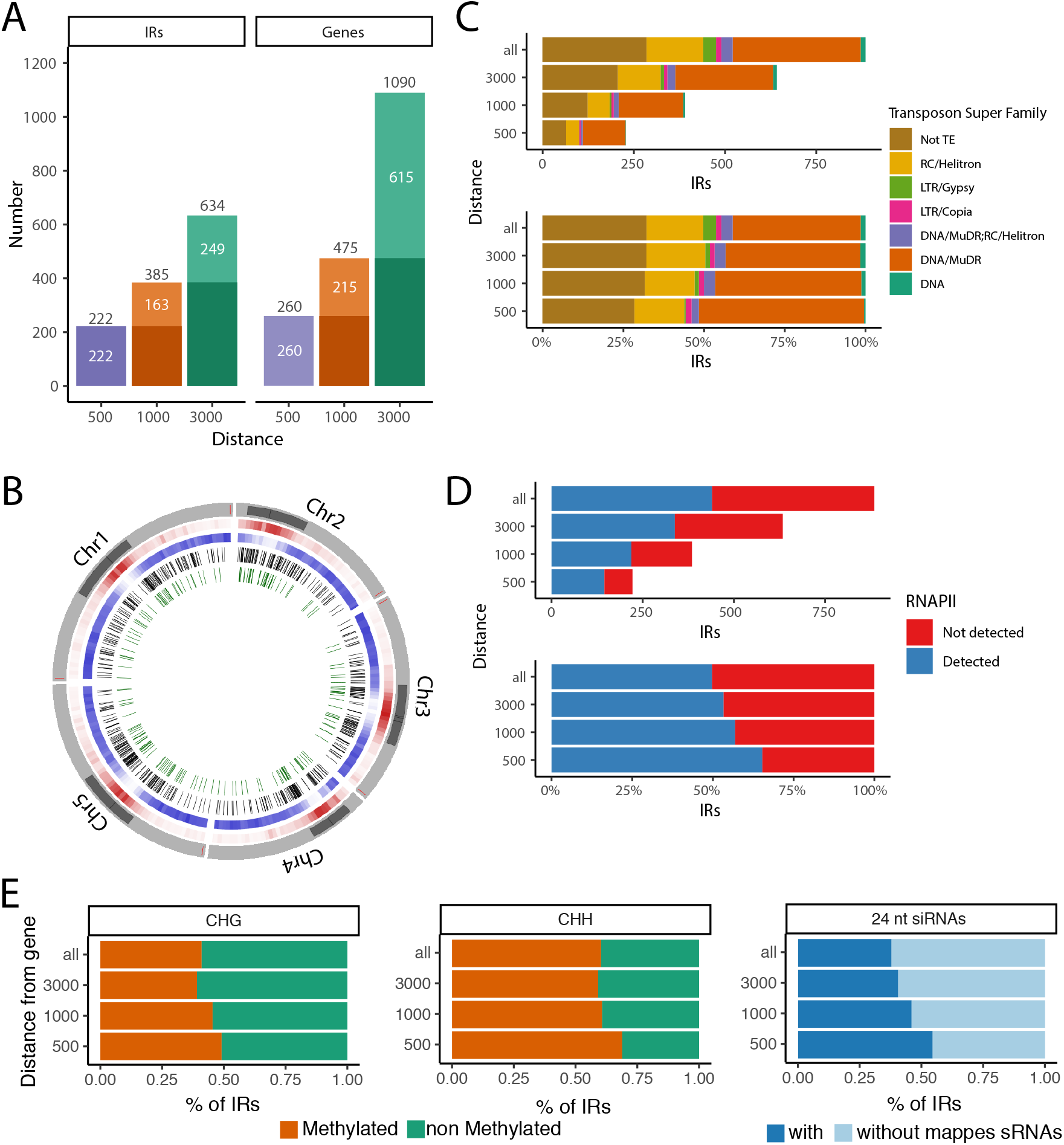
IRs distribution near annotated genes in the *A. thaliana* Col-0 reference genome. **A**. Number of IRs detected in the Col-0 *genome* within 500, 1,000 and 3,000 bp from annotated genes (left), and the number of associated genes within the same size window from the identified IRs (right). White numbers indicate the number of IR and genes exclusive of the 0-500, 501-1000, and 1001-3000 bp windows while black numbers show the cumulative count. **B**. Distribution of IRs. Outermost track, the five Arabidopsis chromosomes with the pericentromeric regions highlighted in darker gray. Then, TE density in 500 kb windows shown in red, and gene density in the same bins, in blue. In both red and blue tracks, darker color indicates higher density. The inner two tracks show the distribution of all IRs (black bars) or only IRs within 3,000 bp upstream or downstream of annotated protein coding genes (green bars). **C**. Classification of the identified IRs between annotated TE super families, only showing super families with at least five overlapping IRs. The “DNA/MuDR;RC/Helitron” category indicates IRs overlapping with TEs from both superfamilies. The upper panel shows the total number of IR in each superfamily while the bottom panel displays the percentage of each superfamily identified in each distance window from annotated genes. **D**. Total number (up) or percentage (down) of IR nascent transcripts associated with RNAPII as detected by plaNet-Seq (Kindgren *et al*., 2020). IRs producing transcripts associated with RNAPII are marked in blue and IRs without RNAPII transcripts in red. **E**. Percentage of gene-adjacent IRs with cytosine methylation in the CHG (left) and CHH (center) contexts detected in at last 10% of the mapping reads and with 24-nt siRNAs mapping to the IR sequence (at least 10 reads in at least one replicate).

With small RNA (sRNA)- and bisulfite (BS)-sequencing, and paralleling what we observed in the case of the *HaWRKY6* adjacent IR, we found that most of the identified IRs near genes produce RNAPII transcripts (Fig. 1D), giving rise to 24-nt siRNAs (486/634 IRs 3000bp away from genes), and are associated with CHG and CHH methylation (Fig. 1E). We often observed an additional peak of methylation, which could represent a second anchor point for chromatin loop formation, either on the opposite border or inside of many genes located near IRs (Fig. S1), once more providing a scenario similar to that of the *HaWRKY6* locus where two close-by methylated regions served as anchor points for regulatory short-range chromatin loops.

To investigate whether the identified IRs, especially those producing siRNAs and DNA methylation, impact local chromatin organization, we extracted RNA and DNA from Col-0 wild-type plants, triple *dcl2, dcl3, dcl4* (*dcl234*)mutants (which are in impaired in 24-nt siRNA production) (Lu et al., 2006), and triple *drm1, drm2, cmt3* (ddc) mutants (impaired in CHH methylation) (Kurihara et al., 2008), and performed RNA-seq, sRNA-seq, BS-seq, and Capture-C. As the sequencing depth required to detect short-range chromatin interaction through standard Hi-C would be enormous, we selected 290 loci containing IRs and 40 control loci and performed a Capture-C experiment to only focus on these regions and increase the chances to detect local chromatin loops. A test mapping of Capture C reads confirmed the enrichment on the captured regions compared to the input HiC samples (Fig. S2A). Collectively, the designed probes cover ∼1% of the genome. After Capture-C we increased the ratio of reads mapping the targeted regions 40 times in average (Fig. S2B).

sRNA-seq revealed that 486 out of the 634 IRs 3000 bp from genes produce 24 nt siRNAs (Fig. 2A), with siRNA levels severely reduced in both *dcl234* and *ddc* mutants, as expected for this class of small RNAs (Fig. 2A). Both CHG (∼50%) and CHH (∼70%) methylated regions strongly overlapped with IRs within 500 bp from genes (Fig. 1E), and methylation in both contexts is reduced in both in *dcl234* and *ddc* for many IRs as expected by the reduction in siRNAs in these genotypes (Fig. 2B). The proportion of IRs with reduced methylation in the mutants is slightly higher for those within 500 bp of genes compared to the other analyzed distance windows (Fig S2C). Changes in siRNAs and methylation were correlated, consistent with a drop of siRNAs leading to reduced methylation in RdDM impaired mutants (Fig. 2C). Altogether, these data suggest that a large proportion of IRs located near genes and transcribed by RNAPII may be able to trigger DNA methylation in *cis* through the non-canonical RdDM pathway.

**Figure 2.**
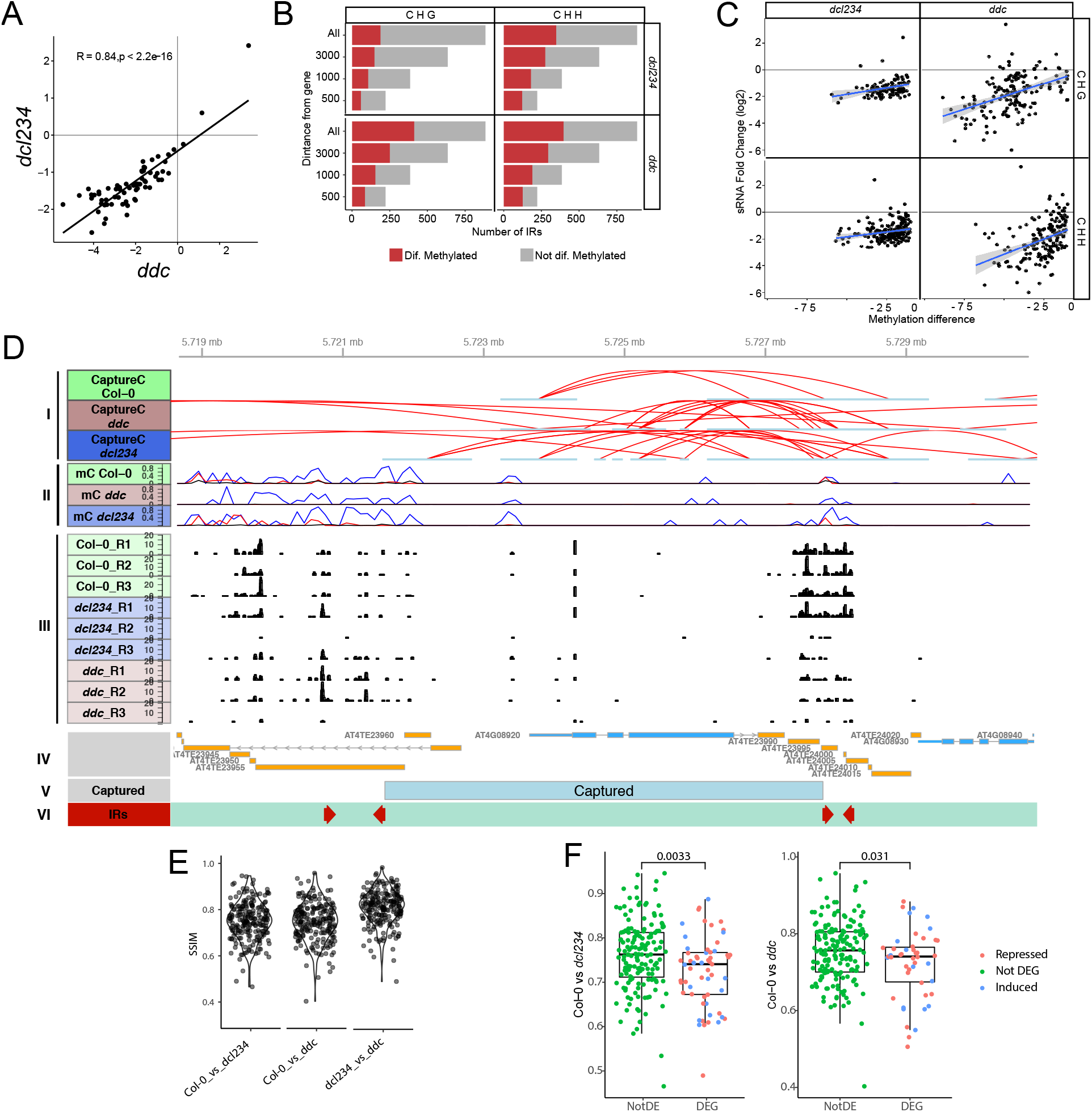
IR near genes produce siRNAs, trigger DNA methylation and alter the local chromatin 3D organization. **A**. Changes in siRNA levels mapping to IRs in the *ddc* and *dcl234* mutants with respect to Col-0. The axes represent log2 Fold Change of siRNA reads per million (RPM) over each IR. The correlation (R) is shown on the upper left corner. **B**. Number of gene-associated IRs showing differential DNA methylation in the CHG and CHH contexts in the *ddc* and *dcl234* mutants with respect to Col-0. **C**. Scatter plots showing the relation between changes in siRNA levels and DNA methylation over each gene-associated IRs in both mutants, *ddc* and *dcl234*. The blue lines show the linear regression while the gray shades show the confidence interval. **D**. Region of the Col-0 genome containing the *CRY1* locus displaying the epigenetic and topological profile. (I) Chromatin interactions as detected by Capture-C experiments and shown by red lines indicating interacting fragments. (II) Cytosine DNA methylation in CG (blue), CHG (red) and CHH (black) contexts. (III) 24 nt siRNAs mapping to the genomic regions as determined by sRNA sequencing in biological triplicates of WT plants and *dcl234* and *ddc* mutants. (IV) Annotated genes (cyan) and transposons (yellow) in the region. (V) Region captured by the probes designed for the CaptureC experiment. (IV) IRs identified in this region of the genome. **E**. Structural similarity (SSIM) of captured regions in *dcl234* and *ddc* mutants with respect to Col-0 or to each other. **F**. SSIM of captured regions in *dcl234* and *ddc* mutants with respect to Col-0 grouped by regions including differentially regulated genes (DEGs) or regions without DEGs (NotDEGs). The colors of the dots represent whether they are not differentially expressed, upregulated, or downregulated.

As in the case of the *HaWRKY6* locus in sunflower, our data suggest that IRs located near genes can act as regulatory elements changing the epigenetic landscape of the region. In order to assess the impact of IR methylation on the surrounding chromatin organization, we used the software CHESS to compare the structural similarity (SSIM) of the IR-host regions between wild type and mutants (Galan et al., 2020). We also statistically determined the specific anchor points of chromatin loops in each sample using capC-Map (Buckle et al., 2019) in combination with peakC (Geeven et al., 2018). Analyzing individual loops, we found clear alterations in the chromatin topology in several randomly picked loci (Fig. 2D and S3). To compare the differential methylation-related changes in loops formed near IRs, we calculated the SSIM for each captured region plus 10 kb on each end in wild type and siRNA- or methylation-deficient mutants. The choice of a global similarity measure, the SSIM, to study short-range chromatin changes, rather than comparing individual loops, aimed at increasing the power to detect reliable differences, as random interactions increase with shorter distances and increase the methodological background noise. Moreover, other interactions, caused by dimerization of DNA-bound transcription factors or nucleosome packing, can also impact such analyses. Chromatin organization of the IR-containing loci, which should have SSIM values ∼1 if similar between the genotypes, were often changed in *ddc* and *dcl234* mutants (Fig. 2E), indicating that (IR-triggered) methylation has a substantial effect on the local 3D topology. Such differences are less pronounced when comparing *ddc* with *dcl234* as could be expected from the methylation deficiency observed in both, therefore lacking the anchor points for loop formation (Fig. 2E).

Changes in chromatin topology can affect gene expression (Domb *et al*., 2022). Given the alterations in chromatin topology that we found, we wondered whether they impact gene expression. Formation of a chromatin loops may repress or activate gene transcription, or even trigger production of alternative transcripts (Gagliardi and Manavella, 2020).

We found 4,305 differentially expressed genes (DEGs) in the *ddc* mutant in comparison with Col-0, and 3,636 DEGs in *dcl234*, many of them overlapping between the two mutants (Fig. S4A). When we compared the SSIMs in regions with DEGs with SSIMs in regions without DEGs we found topological differences (lower SSIM) to be significantly increased in regions that included DEGs (Fig. 2F). We then split those differences into regions linked to differentially and not differentially methylated IRs, which revealed greater topological differences in *ddc* mutants for regions both including DEGs and differentially methylated IRs (Fig. S4B). Methylation seemed to affect the SSIM correlation with DEGs less clearly in *dcl234* (Fig. S4C). Loci with altered topology in *ddc* and *dcl234* included both up- and downregulated genes (Fig. 2F). Thus, changes in chromatin topology caused by an impaired RdDM machinery could be linked to opposite changes in gene expression depending on the loci. This observation, also detected in rice for genes adjacent to MITEs (Lu *et al*., 2012), fits with short-range chromatin loops affecting RNAPII activity in different ways, depending on which part of the gene is included in the loop (Gagliardi and Manavella, 2020).

In summary, our data indicated that the insertion of an IR near a gene can trigger changes in the local chromatin topology that ultimately affect gene expression in a locus- and loop-dependent way.

### The absence of IRs near genes between Arabidopsis accessions causes natural variation in the local chromatin topology

TEs are commonly silenced in plants as a means of protecting genome integrity. However, under extreme stress conditions TE transcription can be reactivated, giving TEs the potential to jump in the genome in a process that has been proposed to help adaptation to new environments (Ito et al., 2011; Tittel-Elmer *et al*., 2010). We wondered whether the IRs we were studying could have adaptative potential to change the structure of the RNAs produced by a locus or their expression, through changes in the 3D chromatin organization.

To test this hypothesis, we first used published TE polymorphism datasets (Stuart et al., 2016) to detect variations in the TE content of 216 *A. thaliana* genomes (Schmitz et al., 2013). Narrowing this down further to the 634 IRs located within 3,000 bp upstream or downstream of annotated genes in the Col-0 reference genome, we found 193 of these 634 IRs to be variable between the 216 accessions studied (Fig. 3A). IRs without a clear TE origin are underrepresented in the collection of polymorphic IRs (Fig. 3A). These 193 IRs are located within 3,000 bp of 368 annotated genes. This implies that each IR can influence two or more adjacent genes in many cases, an observation that is not surprising given the compact nature of *A. thaliana* genomes. To investigate the effect that the variation in the IR content may have on local chromatin topology and gene expression, we repeated the Capture-C, RNA-seq, BS-seq, and sRNA-seq analyses in the Ba1 and Hod accessions. We selected these two accessions based on the number of variable IRs present in each of them (46 and 31 IRs present in Col-0 are missing in Hod and Ba-1, respectively, with 18 missing in both accessions). With the Capture-C experiment, we captured 290 regions comprising the genomic sequences around variable IRs which included adjacent genes. The analysis of individual loci revealed that the presence of an IR at a locus correlated with the accumulation of 24 nt siRNAs and with distinctive short-range chromatin loops (Fig. 3B and S3).

**Figure 3.**
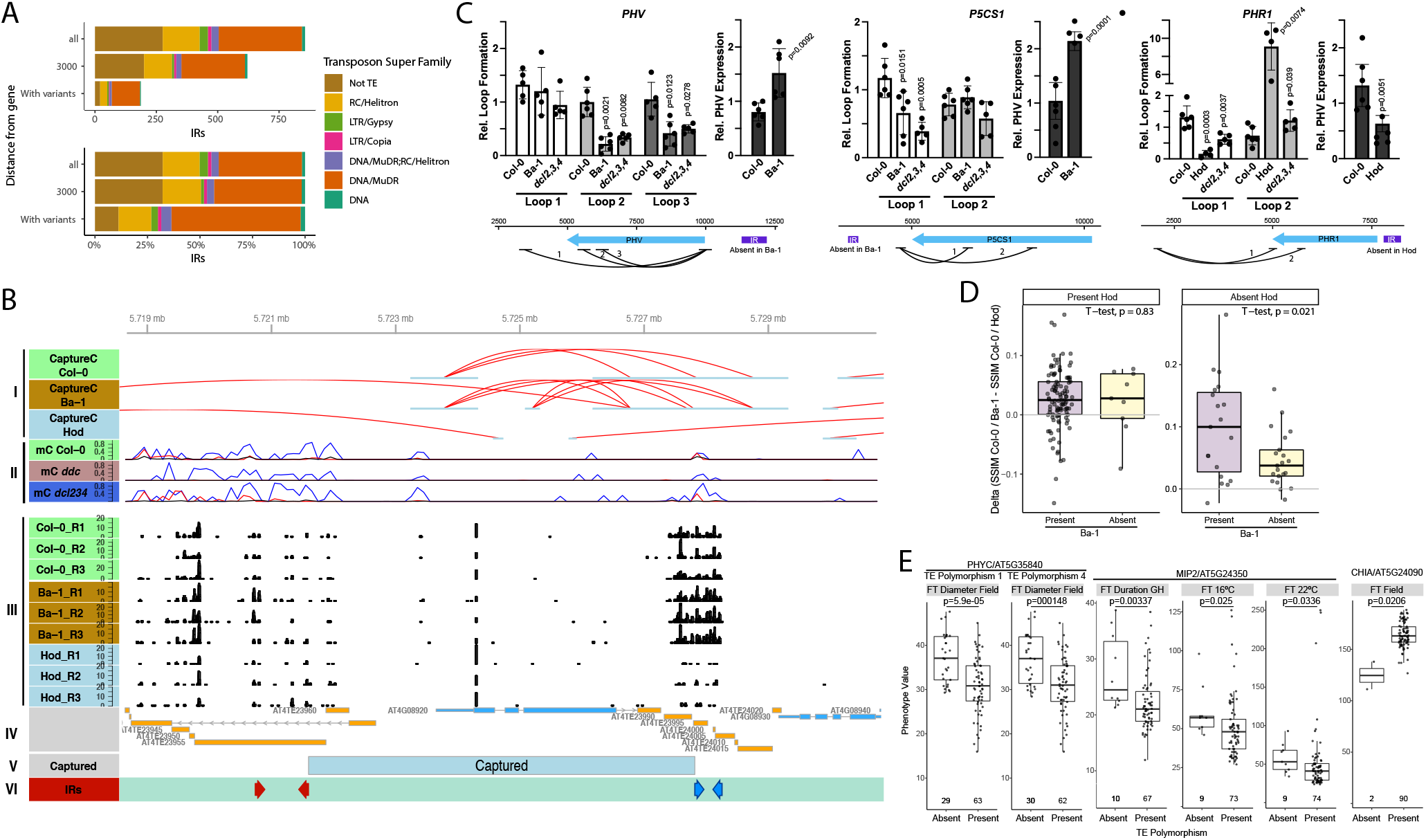
Insertional polymorphisms of IRs near genes cause natural variation in short-range chromatin topology. **A**. Number of IRs and polymorphic IRs associated with genes (upper panel). Percentage of IRs and polymorphic IRs by annotated TE superfamilies (lower panel). Only superfamilies with at least five overlapping IRs are displayed. The “DNA/MuDR;RC/Helitron” category indicates IRs overlapping with TEs from both superfamilies. **B**. Region of the Col-0 genome containing the *CRY1* locus with its epigenetic and topological profile. (I) Chromatin interactions as detected by Capture-C experiments from Col-0 and Ba-1 plants (both contain the IR) and Hod plants (which has a deletion polymorphism in the same IR). Red lines indicate interacting fragments. (II) Cytosine DNA methylation in CG (blue), CHG (red) and CHH (black) contexts. (III) 24 nt siRNAs mapping to the genomic regions as determined by sRNA sequencing in biological triplicates of wild-type Col-0, Ba-1 and Hod plants. (IV) Annotated genes (cyan) and transposons (yellow) in the region. (V) Region captured by the probes designed for the Capture-C experiment. (IV) IRs identified in this region of the genome. The IR is red is conserved between the three genotypes while the IR in blue is missing in Hod. **C**. RT-qPCR with total RNA, and Chromatin Conformation Capture (3C) qPCT experiments to quantify the formation of specific chromatin loops as well as the expression of the affected loci in WT, Ba-1, Hod plants and *dcl234* and *ddc* mutants. Data are presented as mean values +/- SD. p-values were calculated with two-tailed unpaired Student’s t-Test with Welch’s correction. n>4 biologically independent samples. A diagram on the bottom of each gene shows the chromatin loops amplified and confirmed by Sanger sequencing for each locus. **D**. Delta of the structural similarity (SSIM) calculated with CHESS for the contrasts Col-0 vs Ba-1 and Col-0 vs Hod for each captured region. Each captured region is classified depending on whether the associated IR is present or absent in the three analyzed accessions. **E**. Association of flowering phenotypes (FT), including rosette diameter upon flowering, flowering duration, and flowering time at different temperatures either in plants grown in the field or green house (GH), with the presence or absence of an IR near selected genes.

We chose five potential candidate genes (*PHV, PHYC, P5CS1, PHR1, CRY1*) to evaluate the effect of polymorphic IRs on loop formation and gene expression. We corroborated by Sanger sequencing that each of the loci has IRs in Col-0, but not in the indicated accessions (Fig. 4B, 4G, andS5). We then used Chromatin Conformation Capture (3C) followed by qPCR to confirm and quantify the formation of IR-dependent chromatin loops at these loci, and RT-qPCR to measure correlation with gene expression in each locus (Fig. 3C, 4C, and 4I). We detected alternative chromatin loops at *PHV, P5CS1*, and *PHR1* (Fig. 3C). In the case of *PHV*, loop 1 appeared to form independently of the associated IR, but loops 2 and 3 were missing in Ba-1, the accession without the IR, or in *dcl234* mutants (Fig. 3C). For *P5CS1*, loop 2 appeared independently of the presence of the IR while the formation of an intragenic loop 1 correlated with the presence of the IR or a functional RdDM machinery (Fig. 3C). Both for *PHV* and *P5CS1* the absence of a loop-triggering IR appeared to be associated with enhanced gene expression (Fig. 3C). In the case of *PHR1*, we found that the formation of a loop 1 depends on the presence of the IR, which is missing in the Hod accessions, but loop 2 was only formed when the IR is missing, probably reflecting a hierarchy of chromatin interactions controlled by the IR (Fig. 3C). Contrary to the results with *PHV* and *P5CS1*, the absence of the IR near *PHR1* caused repression of the gene.

**Figure 4.**
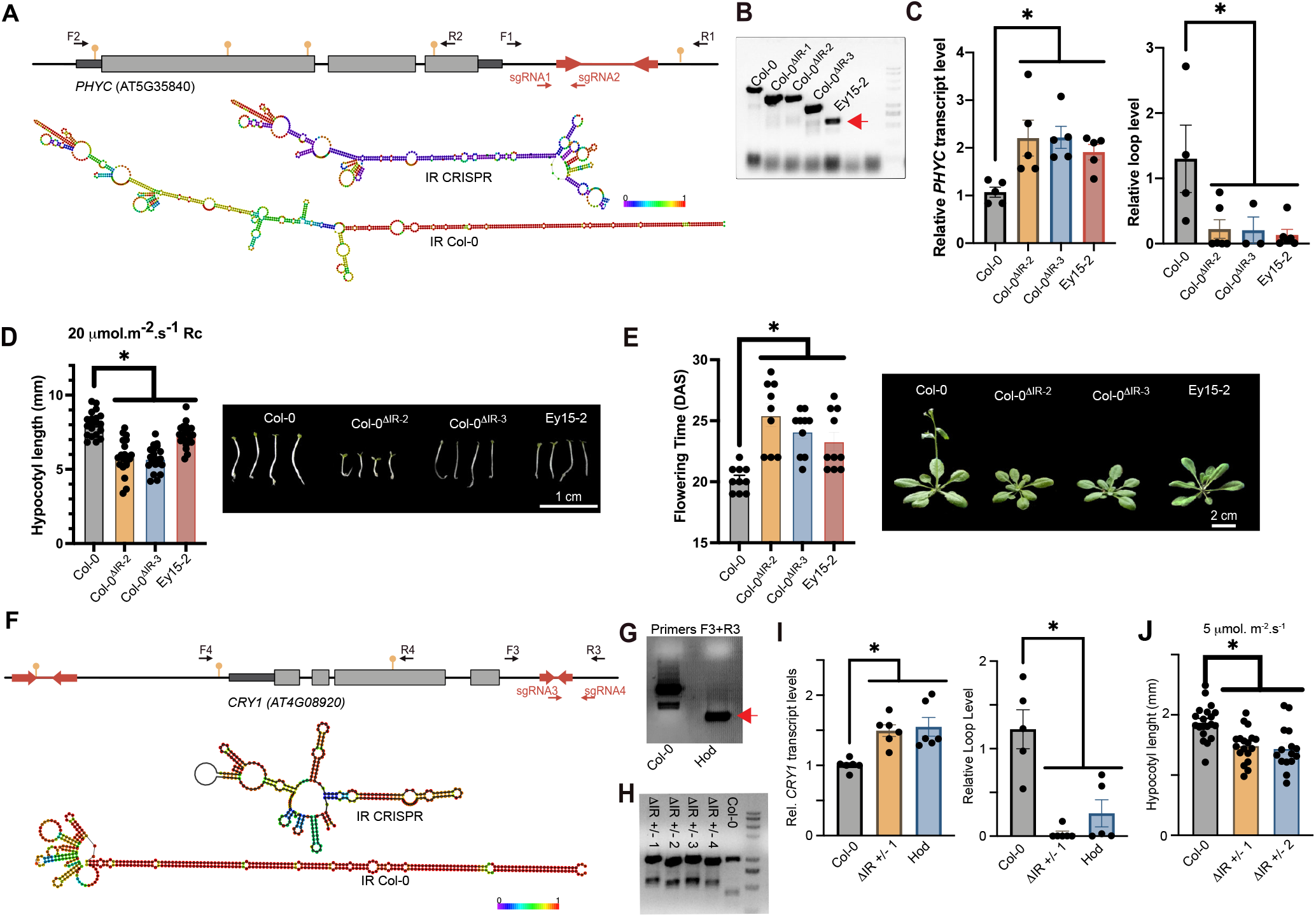
Insertional polymorphisms of IRs near *PHYC* and *CRY1* cause changes in the loci topology, expression and associated phenotypes. **A**. *PHYC* locus diagram showing the position of the IR, restriction sites for EcoRI used for 3C experiments (yellow pins), the sgRNAs used for CRISPR/Cas9 deletion (sgRNA1 and sgRNA2), and primer positions. Predicted secondary RNA structures for the IR transcript in Col-0 plants or in the CRISPR-edited plants are shown using a color scale corresponding to base pairing probability 0 (purple) to 1 (red). **B**. PCR using F1-R1 primers on genomic DNA, confirming partial deletion of the IR in the CRISPR lines and complete absence of the IR in Ey15-2 accession. Red arrowhead indicates the band sequenced to confirm the absence of the polymorphic IR in the Ey15-2 accession. **C**. *PHYC* relative expression as measured by RT-qPCR and normalized against *ACT2* (left panel). Quantification of the formation of a short-range chromatin loop by 3C-qPCR analysis using EcoRI and F2-R2 primers (right panel). Data represents individual values + SEM of five independent biological replicates. **D**. Hypocotyl length under Continuous Red light (Rc) regimen. Seedlings were grown under Rc light 20 μmol m-^2^ s^-1^ for four days before measurement. **E**. Flowering time for the different genotypes expressed as days after sowing (DAS). Data represents individual values ± SD, significant differences are indicated as *, for P<0.05 in a one-way ANOVA. **F**. *CRY1* locus diagram showing the position of the IRs, restriction sites for XbaI (yellow pins), the sgRNAs used for CRISPR/Cas9 deletion (sgRNA3 and sgRNA4), and primer positions. On the bottom are the secondary RNA structures as predicted by RNAfold for the Col-0 wildtype IR transcript and the CRISPR-edited outcome of the same IR. Color scale shows base pair probability 0 (purple) to 1 (red). **G and H**. PCR using F3-R3 primers upon genomic DNA showing successful partial deletion of the IR in CRISPR heterozygous lines and complete absence in Hod plants. Red arrowhead indicates the band sequenced to confirm the absence of the polymorphic IR in the Hod accession. **I**. *CRY1* expression analysis measured by RT-qPCR normalized against *ACT2* on the left. Quantification of a chromatin loop by 3C-qPCR using XbaI and F4-R4 primers (right panel). Data represents average values of three independent biological replicates with SEM. **J**. Hypocotyl length in continuous blue light. Seedlings were grown under blue light 5 μmol m^-2^ s^-1^ for four days before measured. In all cases data represent individual values ± SD, significant differences are indicated as *, for P<0.05 in a one-way ANOVA.

When we analyzed the genome-wide effects of insertional IR polymorphisms over genome topology using CHESS, we also found a drastic change in chromatin folding associated with the presence/absence of an IR (Fig. 3D). For this analysis we calculated the difference between the SSIMs obtained from the comparison of Ba-1 and Hod genome topology against Col-0, as recommended by the authors of CHESS, to have a common reference to score differences between these accessions (Galan *et al*., 2020). As it can be observed in Figure 3D, the difference in SSIMs are greater when Ba-1 shares the IR with Col-0, having a higher SSIM, but it is absent in Hod resulting in a lower SSIM and an increased positive difference.

These data suggest that the insertion of an IR near coding genes can have an impact on local chromatin topology that affects gene expression. Aiming to explore whether this could potentially have adaptive consequences, we asked whether there is a correlation between the presence/absence of an IR near a gene and well-defined phenotypic traits recorded in the Arapheno database (Seren et al., 2017). This analysis revealed an association between IR insertional polymorphisms near some genes with a given phenotype related to flowering, a typically adaptative trait (Fig. 3E). This observation suggests that insertion of an IR near a gene can not only impact the local chromatin topology, but that it may also have adaptive implications by changing the phenotypes controlled by the adjacent gene.

### The insertion of an IR near genes causes strong phenotypic effects

Our data suggest that the insertion of an IR near a gene can be a significant event during adaptative evolution. However, this association may not be caused by the IR itself, but by adjacent polymorphisms that may be in linkage disequilibrium with the IR. In addition, even if the association is with the IR itself, it may not involve changes in the genome topology. To study this possibility we selected two loci, *PHYC* and *CRY1*, which display natural variation in the presence/absence of a close-by IR that is associated with developmental phenotypes (Fig. 3E). Both *PHYC* and *CRY1* associated IRs are located downstream of the transcription termination site (TTS) (Fig. 4A and 4F), thus making it less likely that they act directly on the promoter of the genes as general regulatory elements.

*PHYC* encodes a photoreceptor capable of sensing red light (R) and far-red light (FR) and is implicated in several developmental transitions, such as flowering, seed germination and hypocotyl elongation (Chen et al., 2014; Kippes et al., 2020; Li et al., 2020; Nishida et al., 2013). In Col-0 an IR is located ∼500 bp downstream of the TTS of *PHYC*, while it is missing in several natural accessions, including, for example, Ey15-2 (Fig. 4A and 4B). To test whether this IR impacts *PHYC* expression and related phenotypes, we used a CRISPR/Cas9 strategy to delete a fragment of the IR, with which we can disrupt dsRNA formation without completely removing the IR sequence (Fig. 4A). Three homozygous lines were obtained with a deletion of the IR fragment (Fig. 4B). Supporting a role for this IR in modulating genome topology and gene expression, we detected a chromatin loop encompassing the entire *PHYC* gene in Col-0 plants that was absent in the CRISPR mutant lines and in the Ey15-2 accession, which lacks the IR (Fig. 4C). The absence of the IR, both in Ey15-2 and in the CRISPR lines, correlated with higher expression of *PHYC*, indicating that the IR promotes the formation of a repressive loop in Col-0 (Fig. 4C). Both, the CRISPR lines and Ey15-2, had altered developmental responses related to known *PHYC* functions, including delayed flowering and shortened hypocotyls under a continuous red-light treatment (Fig. 4D and 4E). Altogether, our data indicate that the presence of the IR next to *PHYC* has a substantial impact on gene regulation, by enabling the formation of a short-range chromatin loop that represses gene expression, thereby contributing to the differential response of natural accessions to light signals.

CRY1 is a blue light receptor and participates predominately in the regulation of blue-light inhibition of hypocotyl elongation and anthocyanin production (Ahmad et al., 1998; Ahmad et al., 1995; Liu et al., 2022). Two IRs are located near the gene, ∼2,000 bp upstream of the transcription start site (TSS) and ∼550 bp downstream of the TTS (Fig. 4F). The second IR is variable among *A. thaliana* accession, missing, for example, in Hod (Fig. 4G). Using CRISPR/Cas9 engineering, heterozygous lines could be obtained missing a fragment of the IR located downstream of the TSS (with homozygous lines apparently not being viable) (Fig. 4H). Similar to *PHYC*, we detected a chromatin loop that brings together the borders of the *CRY1* gene only in those plants containing the IR and that represses the expression of the gene (Fig. 4I). The absence of the IR, both in the CRISPR lines or in the natural accession Hod, correlated with higher expression levels of *CRY1* and shorter hypocotyls under blue light (Fig. 4I and 4J), a phenotype previously described in *CRY1* overexpressing lines (He et al., 2019; Liu *et al*., 2022).

## Discussion

How organisms can adapt to a rapidly changing environment is one of the most interesting questions in evolutionary biology (Barrett and Schluter, 2008; Hermisson and Pennings, 2005). SNPs have been the main focus of many genomic studies aiming to assess the evolutionary potential of mutations, but TEs can be particularly powerful actors in rapid adaptation, as single transposition events can have potentially wide-ranging consequences on gene expression and derived phenotypes (Baduel *et al*., 2021). On one side, the broad distribution of TEs across the genome facilitates the generation of chromosomal rearrangements through ectopic recombination. Even more significantly, the mobilization of TEs can disrupt, modify or even change the expression of genes in various ways that could generate a favorable adaptative trait (Dubin *et al*., 2018; Friedli and Trono, 2015). New alleles caused by TE insertions have been proposed to guarantee a consistent supply of potentially adaptable variants in response to the environment (Baduel *et al*., 2021). Still, many TEs inserted within gene-rich areas are quickly purged, according to population genomic surveys of TE polymorphisms (Quadrana et al., 2016). This is in agreement with transposition tending to produce alleles with negative effects.

Contrary to autonomous TEs, MITEs were found to be distributed on chromosome arms in plants, highly associated with genes, and frequently transcribed with adjacent genes (Kuang et al., 2009; Lu *et al*., 2012; Oki et al., 2008). In agreement with this, our study identified numerous TE-derived IRs near coding genes in *A. thaliana*. Different from rice, where more than half of the genes are associated with MITEs (Lu *et al*., 2012), fewer genes are associated with IRs in *A. thaliana*. This observation is not surprising as Arabidopsis is an outlier regarding TE content among plants, with only 15% of its genome represented by TEs. In comparison, they account for 85% of maize and up to ∼40% of rice genomes (Arabidopsis Genome, 2000; Li et al., 2017; Schnable *et al*., 2009).

The association between MITEs and protein coding genes suggested that these TEs may play essential roles in genome evolution. Current evidence suggests that siRNA-triggered TE methylation tends to cause the repression of neighboring genes (Hollister and Gaut, 2009; Hollister et al., 2011). However, in the case of MITEs, both positive and negative effects on the expression of host genes have been reported (Gagliardi *et al*., 2019; Underwood *et al*., 2022; Wu *et al*., 2022; Xu *et al*., 2020; Zhang *et al*., 2016). The weak correlation between methylation of MITEs and the expression of adjacent genes can now be better explained in the light of our findings showing that IRs located near genes affect the local chromatin organization. While methylation of DNA on its own is expected to have primarily repressive effects, short-range chromatin loops could produce many different outcomes, including transcriptional repression/activation and production of alternative mRNAs (Gagliardi and Manavella, 2020).

MITEs have the potential to transpose into various locations in the genome resulting in the presence/absence (insertional) polymorphisms between genotypes (Lu *et al*., 2012; Lyons et al., 2008). Such polymorphism can be caused by the insertion or excision of a MITE from a locus. However, it is unknown which of these scenarios contributes more to the genetic variation within a species. Here, we show that ∼30% of the IRs located within 3,000bp of protein coding genes present insertional polymorphism between 216 *A. thaliana* natural accessions. Our results also indicate that such natural variation of the gene-associated IR content can cause changes in chromatin organization that could be considered “3D polymorphisms”. Our experiments using not only association between IR polymorphisms and gene expression changes, but also CRISPR/Cas9-editing to demonstrate a causal relationship in several cases, show that the insertion of an IR near a gene can have a profound impact on chromatin organization, gene expression, and associated phenotypes. This phenomenon can potentially boost a plant’s capacity to adapt to a rapidly changing environment. Because IRs can coopt the promoters of adjacent genes to produce RNAPII-derived siRNAs through a stem-loop dsRNA intermediate, they can act as autonomous regulatory elements, drastically changing the chromatin landscape of a locus upon insertion. These characteristics turn IRs into powerful elements during adaptative evolution.

Modelling has suggested faster generation of large-effect alleles due to larger transposition rates in specific populations in response to global warming (Baduel *et al*., 2021). Consistent with this scenario, and in a world of a rapidly changing climate, the discovery of IRs as elements shaping the 3D chromatin organization and driving genome adaptation is of great interest. The manipulation of IRs, and in consequence, genome topology, can potentially become a powerful biotechnological tool to improve crop adaptation without the need to incorporate exogenous DNA or alter coding sequences in plants.

## Material and methods

### Plant material and growth conditions

*Arabidopsis thaliana* accessions Col 0, Hod, Ba1, EY15-2, *dcl234* (Lu *et al*., 2006), and *ddc* (Kurihara *et al*., 2008) mutants and CRISPR mutant lines were grown at 23ºC in long day (16/8 hours light/dark). Blue and red-light experiments were carried out as follows: seeds were sowed on petri dishes with humidified filter paper and stratified in the dark for 5 days, and then transferred to white light (80 μmol m^-2^ s^-1^) at 23ºC for 3 hours and subsequently transferred to red (20 μmol m^-2^ s^-1^) or blue light (5 μmol m^-2^ s^-1^) in LED chambers. Hypocotyl measurements were taken at four days using ImageJ (http://rsb.info.nih.gov/ij).

To obtain plants with genomic fragments deletions a CRISPR/Cas9 vector toolbox (Wu et al., 2018) was used. Specific sgRNAs, described in Supplemental Table S1, were designed to obtain the IRs deletions. Col-0 plants were transformed using the floral dip method, T1 plants were selected based on the presence of red fluorescence in seeds under a fluorescent dissecting microscope (Leica, Solms, Germany), non-fluorescent T2 seeds missing the transgenes were grown and genotyped by PCR to identify effective deletions.

For genomic DNA extraction 100 mg of fresh plant material was ground in 700 µl of extraction buffer (200 mM Tris-HCl pH8, 25 0mM EDTA, 0.5% SDS) and precipitated with isopropanol. PCR was performed using primers detailed in Table S1. RNA was extracted using TRIzol™ Reagent (Invitrogen) following manufacturer’s recommendations.

The presence of the IR in different *A. thaliana* accessions was determine by amplification of the IR region with flanking primers with Q5 polymerase, followed by Sanger sequencing. For the deletion of the IRs neighboring *PHYC* and *CRY1*, pairs of sgRNAs targeting each locus were cloned in a CRISPR/Cas9 super module (SM) vector as described (Wu *et al*., 2018). Briefly, sgRNAs targeting flanking regions of the IR were designed using the CRISPR-P Web Tool (Lei et al., 2014). Each sgRNA was introduced into the shuffle vectors by overlap PCR with Q5 Hi-Fidelity polymerase followed by digestion of the original vector with *DpnI* (Thermo Scientific). A destination vector harboring *UBQ10* promoter, pcoCas9, proLacZ:LacZ between both sgRNAs targeting each IR, and At2S3:mCherry for fluorescence selection in seeds were generated with the Green Gate assembly system. Destination vectors were transformed into Col-0 plants, and red fluorescence-positive seeds were isolated as hemizygous seeds. Transgene-free T2 offspring without seed fluorescence were chosen, and plants were tested by PCR and Sanger sequencing to identify IR deletion lines. All primers used are listed in Table S1

### IR detection

Inverted repeats were identified in the Col-0 *Arabidopsis thaliana* genome using einverted, from the EMBOSS program suite (Rice *et al*., 2000). The paremeters used were: maximum repeat of 1000, a gap penality of 8, a minimum score threshold of 150, a match score of 3, and a mismatch score of -4.

### RNA-seq

RNA-seq library preparation was performed as described (Cambiagno et al., 2021). An in-house scaled-down version of Illumina’s TruSeq reaction was used. mRNA was purified with NEBNext Poly(A) Magnetic Isolation Module (New England Biolabs, Ipswich, MA) and heat fragmented with Elute-Prime-Fragment buffer (5x first-strand buffer, 50 ng/ml random primers). For first- and second-strand synthesis SuperScript II Reverse Transcriptase (Thermo Fisher) and DNA polymerase I (NEB) were used, respectively. T4 DNA polymerase, Klenow DNA polymerase, T4 polynucleotide kinase (NEB), and Klenow Fragment (30/50 exo-) (NEB) were used for end repair and A-tailing. Ligation of universal adapters compatible with Nextera barcodes i7 and i5 was performed with T4 DNA Ligase (NEB), and Q5 Polymerase (NEB) was used for PCR enrichment using Nextera i7 and i5 barcodes. SPRI beads were used for DNA purification in each step and size selection of the library preps. 2 × 150 bp paired-end reads were generation on the Illumina HiSeq 3000 platform.

The analysis started by quality trimming and filtering the raw reads with Trimmomatic version 0.36; (Bolger et al., 2014. They were then aligned to the Arabidopsis thaliana genome (TAIR10) using STAR version 2.5.2b, {Dobin, 2013 #73), which was guided by the gene and exon annotation from Araport V11 201606, (Pasha et al., 2020). Samtools version 1.; (Li et al., 2009) was then used to keep only primary alignments with a minimum MAPQ of 3. Read quality before and after trimming was analyzed with FastQC (version 0.11.5; https://www.bioinformatics.babraham.ac.uk/projects/fastqc/) and, together with mapping efficiency, they were summarized with MultiQC version 1.7 (Ewels et al., 2016). Read counts on each gene were then calculated with featureCounts version 1.6.2 (Liao et al., 2014). This pipeline was run with the aid of the Snakemake workflow engine (Koster and Rahmann, 2012). Gene counts were used to identify differentially expressed genes with DESeq2 (Love et al., 2014); R Core Team 2022) filtering out genes with counts below 10 in all samples.

### sRNA-seq

For sRNA-seq library preparation, 1 µg of total RNA was used as input for the TruSeq sRNA Library Preparation kit (Illumina) as described in the TruSeq RNA Sample Preparation v2 Guide (Illumina). BluePippin System (Sage Science) was used for sRNA library size selections. Sequencing was performed on the Illumina HiSeq 3000 platform.

The small RNA reads generated were first cut to remove 3’ adapters using cutadapt (version 1.9.1) and their quality checked using FastQC (version 0.11.4, https://www.bioinformatics.babraham.ac.uk/projects/fastqc/) and MultiQC (Ewels *et al*., 2016). They were then mapped with bowtie (version 1.1.2; (Langmead et al., 2009)) to *A. thaliana* rRNA, tRNA, snoRNA and snRNA from RFAM (version 14.1, (Kalvari et al., 2018). Unmapped reads were then mapped, also with bowtie, to the *A. thaliana* genome. Statistical analyses were performed in the R statistical programming environment (R Core Team, 2022) and graphics were produced with the ggplot package.

IRs which had 10 or more 24 nucleotide reads in its entire region were considered to have potential sRNA production. Changes in sRNA levels were calculated first calculating reads per million (RPM) mapping reads to the genome in each library, then averaging this value for all replicates in each IR, and then calculating the log2 Fold Change between the RPM in the mutant versus Col-0.

### Bisulfite treatment of DNA and library preparation

For BS-seq, DNA extraction was performed with DNeasy plant Mini Kit (QUIAGEN). DNA was sheared to 350 bp by g by Covaris ultrasonication. Libraries were generated with Illumina TrueSeq DNA Nano Kit. After adaptor ligation, libraries were bisulfite converted with the Epitec Plus DNA Bisulfite Conversion Kit (QUIAGEN). Library enrichment was done using Kapa Hifi Uracil+ DNA polymerase (Kapa Biosystems, USA). Paired-end reads (2 × 150 bp) were generation on the HiSeq 3000 platform (Illumina).

The analysis of these reads started by quality trimming and filtering them with Trimmomatic (version 0.36; (Bolger *et al*., 2014)). Then we used the Bismark program (Krueger and Andrews, 2011) to perform the mapping of the reads to the *A. thaliana* Col-0 genome, internally done with Bowtie2 (Langmead and Salzberg, 2012), the deduplication of the alignments and the extraction of the methylation results in the three contexts: CG, CHG and CHH. This output was then analyzed in R (R core team, 2022) with the methylKit package (Akalin et al., 2012). Only Citosines with at least 4 reads were considered, and each sample was segmented with methSeg and methylation levels were calculated for those including at least 4 Cs. For Col-0, segments were collapsed for replicates using the mergeGRangesData function from the BRGenomics package (https://rdrr.io/bioc/BRGenomics/) and IRs with repeats overlapping segments with more than 10 or more percent of CHG or CHH methylation were considered methylated.

Differential methylation in the *ddc* and *dcl234* mutants was also calculated with the mehtylKit package. First replicates were combined with the unite function and then differential methylation calculated with the calculateDiffMeth function, correcting for overdispertion with the MN method, using a q-value threshold of 0.1 and a differential threshold of 15 %. Then IRs with repeats overlapping any of these differential segments was considered differentially methylated.

### Capture-C assay

For Capture-C, Hi-C was performed as described (Liu, 2017). Briefly, we collected 1.5 g of plant tissue, and fixed them with 1% formaldehyde. Nuclei were isolated and finally washed with NEB buffer #3. Nuclei penetration was done by resuspending the pellet in 150 μl 0.5% SDS and incubating them at 62 ºC for 5 min. After that, 435 ul of water and 75 ul of 10% Triton X-100 were added and incubated 37 ºC for 15 min. NEB buffer #3 was added to 1X, and 50 U of DpnII to digest the chromatin over night at 37°C. Incubating the digested chromatin with 10 U Klenow, dTTP, dATP, dGTP, and biotin-14-dCTP at 37°C for 2 h, cohesive ends were filled. Blunt-end ligation of chromatin was performed by adding blunt-end ligation buffer to 1X and 20 U of T4 DNA ligase at room temperature for 4 h. Nuclei were lysed with SDS buffer (50 mM Tris-HCl, 1% SDS, 10 mM EDTA, pH 8.0) and incubated with 10 μg proteinase K at 55°C for 30 min. To reverse the crosslinking, NaCl was added to reach 0.2 M and the samples were incubated at 65 °C overnight. Hi-C DNA was purified by Phenol-Chloroform-IAA method and RNAse A treated. Hi-C DNA was sheared to 500 bp with a Covaris E220 sonicator. DNA was purified and size selected (longer than 300 bp) using Ampure beads. Unligated biotin was removed in a reaction with 0.1 mM dTTP, 0.1 mM dATP and 5 U T4 DNA polymerase incubated at 20°C for 30 min. DNA was purified with Ampure beads and end-repair and adaptor ligation were performed with the NEBNext® Ultra™ II DNA Library Prep Kit by following the manufacturer’s instructions. Biotin affinity purification was then performed by using Dynabeads MyOne Streptavidin C1 beads (Invitrogen). Library amplification was done with Ultra II Q5 Master Mix with universal and selected index primers.

For the Capture step, hybridization capture was performed with the MyBaits system (Arbor Biosciences) following the manufacturer’s instructions. Baits of 80 nucleotides were designed on each end of the digestion fragments corresponding to the captured regions. These regions included all genes within 3000 bp of the IR and the spacer region up to the IR, excluding it. When a region was surrounded by 2 IRs, it was considered a single captured region.

Finally, Capture-C DNA was pared-end sequenced (2 × 150 bp reads) on an Illumina HiSeq 3000. The resulting reads were processed with capC-MAP (Buckle *et al*., 2019), which performs the *in silico* genome digestion, read alignment, the pile-up of interactions, and can generate normalized, binned and smoothed profiles of interaction for each target. For this the Col-0 genome was used, an exclusion zone of 500 bp, a bin size of 500 bp and a step of 250 bp. The results were then processed with the R package peakC (Geeven *et al*., 2018) to determine statistically significant interactions. These loops were visualized with the aid of the R package and Gviz (Hahne and Ivanek, 2016).

For the SSIM calculation, raw pileups were first normalized with FAN-C (Kruse et al., 2020), using the VC-SQRT method on 1000 bp bins. The SSIM value was obtained on a region comprising the captured region extended by 10 kb on both extremes, and using a relative window size of 0.1.

### Data processing, plotting and statistical analysis

Data obtained in the different analysis of the sequencing experiments was further processed and statistically analyzed in R (R Core Team, 2022) using a diversity of packages. Genomic information was handled using GenomicRanges (Lawrence et al., 2013), Biostrings (https://bioconductor.org/packages/Biostrings) GenomicInteractions (Harmston et al., 2015) and rtracklayer (Lawrence et al., 2009) packages.

Plots summarizing information were mostly performed with the ggplot and ggpubr packages. Plots of genomic regions were produced with the Gviz (Hahne and Ivanek, 2016) package. Circular plots were generated with ciclize (Gu et al., 2014).

### Data availability

All sequencing data were deposited at the European Nucleotide Archive (https://www.ebi.ac.uk/ena/) public repository with accession PRJEB53956.

### 3C assay and RT-PCR

3C assay was performed as described (Gagliardi *et al*., 2019). For detection of loops at *PHV* and *PHYC*, EcoRI (NEB) overnight digestion was performed; for *P5CS1, CRY1*, and *PHR1*, XbaI (NEB) was used. For DNA ligation, 100 U of highly concentrated T4 DNA ligase (Thermo) were used at 22°C for 5 h in a 4 mL volume. Reverse crosslinking and proteinase K treatment (Invitrogen) were performed, and phenol/chloroform method was used for DNA purification. For interaction frequency measurement, qPCR was performed using *ACTIN2* as housekeeping gene. All primers used are listed in Table S1.

For quantitative RT-PCR, 1 μg of total RNA was used for reverse transcription reactions using RevertAid RT Reverse Transcription Kit (Thermo Fisher Scientific). qPCR was performed using SYBR green (Thermo Scientific Maxima SYBR Green qPCR Master Mix (2x)). Three biological replicates were used to calculate the standard error of the mean. Standard error of the mean (SEM) was calculated using propagation of error of the 2^-ΔΔCt^ values and expressed in figures as two times the SEM. Statistical significance was tested using a two-tailed, unpaired Student’s t-test. All primers used are listed in Table S1.

## Author contribution

A.L.A. performed the majority of the analyses. D.A.C. prepared the libraries for sequencing and Capture-C experiments. P.L.L. and H.A.B. helped with the design of the Capture-C probes. P.L.L. performed the Capture experiment. R.M. and D.A.C validated the chromatin loops formations and created the CRISPR/CAS9 mutant lines. R.M. characterized the CRISPR/Cas9 mutant lines and performed validation experiments. A.L.A., D.A.C., P.A.M., and D.W. conceived this study; P.A.M and D.W. supervised the work and secured project funding; A.L.A., R.M., D.W., and P.A.M. wrote the manuscript.

## Acknowledgments

This work was supported by grants from ANPCyT (Agencia Nacional de Promoción Científica y Tecnológica, Argentina) and Universidad Nacional del Litoral (UNL) to P.A.M and the Max Planck Society to D.W. P.A.M., A.L.A. and D.A.C are members of CONICET; R.M is a fellow of the same institution. We thank the Deutscher Akademischer Austauschdienst (DAAD) and Company of Biologists for short-term fellowship to D.A.C. and R.M. respectively.

## Footnotes

The authors declare no competing interest.

## References

Ahmad, M., Jarillo, J.A., and Cashmore, A.R. (1998). Chimeric proteins between cry1 and cry2 Arabidopsis blue light photoreceptors indicate overlapping functions and varying protein stability. Plant Cell 10, 197–207. 10.1105/tpc.10.2.197.

Ahmad, M., Lin, C., and Cashmore, A.R. (1995). Mutations throughout an Arabidopsis blue-light photoreceptor impair blue-light-responsive anthocyanin accumulation and inhibition of hypocotyl elongation. Plant J 8, 653–658. 10.1046/j.1365-313x.1995.08050653.x.

Akalin, A., Kormaksson, M., Li, S., Garrett-Bakelman, F.E., Figueroa, M.E., Melnick, A., and Mason, C.E. (2012). methylKit: a comprehensive R package for the analysis of genome-wide DNA methylation profiles. Genome Biol 13, R87. 10.1186/gb-2012-13-10-r87.

Arabidopsis Genome, I. (2000). Analysis of the genome sequence of the flowering plant Arabidopsis thaliana. Nature 408, 796–815. 10.1038/35048692.

Ariel, F.D., and Manavella, P.A. (2021). When junk DNA turns functional: transposon-derived non-coding RNAs in plants. J Exp Bot 72, 4132–4143. 10.1093/jxb/erab073.

Baduel, P., Leduque, B., Ignace, A., Gy, I., Gil, J., Jr., Loudet, O., Colot, V., and Quadrana, L. (2021). Genetic and environmental modulation of transposition shapes the evolutionary potential of Arabidopsis thaliana. Genome Biol 22, 138. 10.1186/s13059-021-02348-5.

Barrett, R.D., and Schluter, D. (2008). Adaptation from standing genetic variation. Trends Ecol Evol 23, 38–44. 10.1016/j.tree.2007.09.008.

Bolger, A.M., Lohse, M., and Usadel, B. (2014). Trimmomatic: a flexible trimmer for Illumina sequence data. Bioinformatics 30, 2114–2120. 10.1093/bioinformatics/btu170.

Buckle, A., Gilbert, N., Marenduzzo, D., and Brackley, C.A. (2019). capC-MAP: software for analysis of Capture-C data. Bioinformatics 35, 4773–4775. 10.1093/bioinformatics/btz480.

Cambiagno, D.A., Giudicatti, A.J., Arce, A.L., Gagliardi, D., Li, L., Yuan, W., Lundberg, D.S., Weigel, D., and Manavella, P.A. (2021). HASTY modulates miRNA biogenesis by linking pri-miRNA transcription and processing. Mol Plant 14, 426–439. 10.1016/j.molp.2020.12.019.

Chen, A., Li, C., Hu, W., Lau, M.Y., Lin, H., Rockwell, N.C., Martin, S.S., Jernstedt, J.A., Lagarias, J.C., and Dubcovsky, J. (2014). Phytochrome C plays a major role in the acceleration of wheat flowering under long-day photoperiod. Proc Natl Acad Sci U S A 111, 10037–10044. 10.1073/pnas.1409795111.

Collier, L.S., and Largaespada, D.A. (2007). Transposable elements and the dynamic somatic genome. Genome Biol 8 Suppl 1, S5. 10.1186/gb-2007-8-s1-s5.

Crescente, J.M., Zavallo, D., Del Vas, M., Asurmendi, S., Helguera, M., Fernandez, E., and Vanzetti, L.S. (2022). Genome-wide identification of MITE-derived microRNAs and their targets in bread wheat. BMC Genomics 23, 154. 10.1186/s12864-022-08364-4.

Cuerda-Gil, D., and Slotkin, R.K. (2016). Non-canonical RNA-directed DNA methylation. Nat Plants 2, 16163. 10.1038/nplants.2016.163.

Deragon, J.M., Casacuberta, J.M., and Panaud, O. (2008). Plant transposable elements. Genome Dyn 4, 69–82. 10.1159/000126007.

Domb, K., Wang, N., Hummel, G., and Liu, C. (2022). Spatial Features and Functional Implications of Plant 3D Genome Organization. Annu Rev Plant Biol 73, 173–200. 10.1146/annurev-arplant-102720-022810.

Dubin, M.J., Mittelsten Scheid, O., and Becker, C. (2018). Transposons: a blessing curse. Curr Opin Plant Biol 42, 23–29. 10.1016/j.pbi.2018.01.003.

Ewels, P., Magnusson, M., Lundin, S., and Kaller, M. (2016). MultiQC: summarize analysis results for multiple tools and samples in a single report. Bioinformatics 32, 3047–3048. 10.1093/bioinformatics/btw354.

Fattash, I., Rooke, R., Wong, A., Hui, C., Luu, T., Bhardwaj, P., and Yang, G. (2013). Miniature inverted-repeat transposable elements: discovery, distribution, and activity. Genome 56, 475–486. 10.1139/gen-2012-0174.

Friedli, M., and Trono, D. (2015). The developmental control of transposable elements and the evolution of higher species. Annu Rev Cell Dev Biol 31, 429–451. 10.1146/annurev-cellbio-100814-125514.

Gagliardi, D., Cambiagno, D.A., Arce, A.L., Tomassi, A.H., Giacomelli, J.I., Ariel, F.D., and Manavella, P.A. (2019). Dynamic regulation of chromatin topology and transcription by inverted repeat-derived small RNAs in sunflower. Proc Natl Acad Sci U S A 116, 17578–17583. 10.1073/pnas.1903131116.

Gagliardi, D., and Manavella, P.A. (2020). Short-range regulatory chromatin loops in plants. New Phytol 228, 466–471. 10.1111/nph.16632.

Galan, S., Machnik, N., Kruse, K., Diaz, N., Marti-Renom, M.A., and Vaquerizas, J.M. (2020). CHESS enables quantitative comparison of chromatin contact data and automatic feature extraction. Nat Genet 52, 1247–1255. 10.1038/s41588-020-00712-y.

Geeven, G., Teunissen, H., de Laat, W., and de Wit, E. (2018). peakC: a flexible, non-parametric peak calling package for 4C and Capture-C data. Nucleic Acids Res 46, e91. 10.1093/nar/gky443.

Grzechnik, P., Tan-Wong, S.M., and Proudfoot, N.J. (2014). Terminate and make a loop: regulation of transcriptional directionality. Trends Biochem Sci 39, 319–327. 10.1016/j.tibs.2014.05.001.

Gu, Z., Gu, L., Eils, R., Schlesner, M., and Brors, B. (2014). circlize Implements and enhances circular visualization in R. Bioinformatics 30, 2811–2812. 10.1093/bioinformatics/btu393.

Guo, C., Spinelli, M., Ye, C., Li, Q.Q., and Liang, C. (2017). Genome-Wide Comparative Analysis of Miniature Inverted Repeat Transposable Elements in 19 Arabidopsis thaliana Ecotype Accessions. Sci Rep 7, 2634. 10.1038/s41598-017-02855-1.

Hahne, F., and Ivanek, R. (2016). Visualizing Genomic Data Using Gviz and Bioconductor. Methods Mol Biol 1418, 335–351. 10.1007/978-1-4939-3578-9_16.

Harmston, N., Ing-Simmons, E., Perry, M., Baresic, A., and Lenhard, B. (2015). GenomicInteractions: An R/Bioconductor package for manipulating and investigating chromatin interaction data. BMC Genomics 16, 963. 10.1186/s12864-015-2140-x.

He, G., Liu, J., Dong, H., and Sun, J. (2019). The Blue-Light Receptor CRY1 Interacts with BZR1 and BIN2 to Modulate the Phosphorylation and Nuclear Function of BZR1 in Repressing BR Signaling in Arabidopsis. Mol Plant 12, 689–703. 10.1016/j.molp.2019.02.001.

Hermisson, J., and Pennings, P.S. (2005). Soft sweeps: molecular population genetics of adaptation from standing genetic variation. Genetics 169, 2335–2352. 10.1534/genetics.104.036947.

Hollister, J.D., and Gaut, B.S. (2009). Epigenetic silencing of transposable elements: a trade-off between reduced transposition and deleterious effects on neighboring gene expression. Genome Res 19, 1419–1428. 10.1101/gr.091678.109.

Hollister, J.D., Smith, L.M., Guo, Y.L., Ott, F., Weigel, D., and Gaut, B.S. (2011). Transposable elements and small RNAs contribute to gene expression divergence between Arabidopsis thaliana and Arabidopsis lyrata. Proc Natl Acad Sci U S A 108, 2322–2327. 10.1073/pnas.1018222108.

Ito, H., Gaubert, H., Bucher, E., Mirouze, M., Vaillant, I., and Paszkowski, J. (2011). An siRNA pathway prevents transgenerational retrotransposition in plants subjected to stress. Nature 472, 115–119. 10.1038/nature09861.

Kalvari, I., Nawrocki, E.P., Argasinska, J., Quinones-Olvera, N., Finn, R.D., Bateman, A., and Petrov, A.I. (2018). Non-Coding RNA Analysis Using the Rfam Database. Curr Protoc Bioinformatics 62, e51. 10.1002/cpbi.51.

Kindgren, P., Ivanov, M., and Marquardt, S. (2020). Native elongation transcript sequencing reveals temperature dependent dynamics of nascent RNAPII transcription in Arabidopsis. Nucleic Acids Res 48, 2332–2347. 10.1093/nar/gkz1189.

Kippes, N., VanGessel, C., Hamilton, J., Akpinar, A., Budak, H., Dubcovsky, J., and Pearce, S. (2020). Effect of phyB and phyC loss-of-function mutations on the wheat transcriptome under short and long day photoperiods. BMC Plant Biol 20, 297. 10.1186/s12870-020-02506-0.

Koster, J., and Rahmann, S. (2012). Snakemake--a scalable bioinformatics workflow engine. Bioinformatics 28, 2520–2522. 10.1093/bioinformatics/bts480.

Krueger, F., and Andrews, S.R. (2011). Bismark: a flexible aligner and methylation caller for Bisulfite-Seq applications. Bioinformatics 27, 1571–1572. 10.1093/bioinformatics/btr167.

Kruse, K., Hug, C.B., and Vaquerizas, J.M. (2020). FAN-C: a feature-rich framework for the analysis and visualisation of chromosome conformation capture data. Genome Biol 21, 303. 10.1186/s13059-020-02215-9.

Kuang, H., Padmanabhan, C., Li, F., Kamei, A., Bhaskar, P.B., Ouyang, S., Jiang, J., Buell, C.R., and Baker, B. (2009). Identification of miniature inverted-repeat transposable elements (MITEs) and biogenesis of their siRNAs in the Solanaceae: new functional implications for MITEs. Genome Res 19, 42–56. 10.1101/gr.078196.108.

Kurihara, Y., Matsui, A., Kawashima, M., Kaminuma, E., Ishida, J., Morosawa, T., Mochizuki, Y., Kobayashi, N., Toyoda, T., Shinozaki, K., and Seki, M. (2008). Identification of the candidate genes regulated by RNA-directed DNA methylation in Arabidopsis. Biochem Biophys Res Commun 376, 553–557. 10.1016/j.bbrc.2008.09.046.

Langmead, B., and Salzberg, S.L. (2012). Fast gapped-read alignment with Bowtie 2. Nat Methods 9, 357–359. 10.1038/nmeth.1923.

Langmead, B., Trapnell, C., Pop, M., and Salzberg, S.L. (2009). Ultrafast and memory-efficient alignment of short DNA sequences to the human genome. Genome Biol 10, R25. 10.1186/gb-2009-10-3-r25.

Lawrence, M., Gentleman, R., and Carey, V. (2009). rtracklayer: an R package for interfacing with genome browsers. Bioinformatics 25, 1841–1842. 10.1093/bioinformatics/btp328.

Lawrence, M., Huber, W., Pages, H., Aboyoun, P., Carlson, M., Gentleman, R., Morgan, M.T., and Carey, V.J. (2013). Software for computing and annotating genomic ranges. PLoS Comput Biol 9, e1003118. 10.1371/journal.pcbi.1003118.

Lei, Y., Lu, L., Liu, H.Y., Li, S., Xing, F., and Chen, L.L. (2014). CRISPR-P: a web tool for synthetic single-guide RNA design of CRISPR-system in plants. Mol Plant 7, 1494–1496. 10.1093/mp/ssu044.

Li, H., Handsaker, B., Wysoker, A., Fennell, T., Ruan, J., Homer, N., Marth, G., Abecasis, G., Durbin, R., and Genome Project Data Processing, S. (2009). The Sequence Alignment/Map format and SAMtools. Bioinformatics 25, 2078–2079. 10.1093/bioinformatics/btp352.

Li, Q., Wu, G., Zhao, Y., Wang, B., Zhao, B., Kong, D., Wei, H., Chen, C., and Wang, H. (2020). CRISPR/Cas9-mediated knockout and overexpression studies reveal a role of maize phytochrome C in regulating flowering time and plant height. Plant Biotechnol J 18, 2520–2532. 10.1111/pbi.13429.

Li, X., Guo, K., Zhu, X., Chen, P., Li, Y., Xie, G., Wang, L., Wang, Y., Persson, S., and Peng, L. (2017). Domestication of rice has reduced the occurrence of transposable elements within gene coding regions. BMC Genomics 18, 55. 10.1186/s12864-016-3454-z.

Liao, Y., Smyth, G.K., and Shi, W. (2014). featureCounts: an efficient general purpose program for assigning sequence reads to genomic features. Bioinformatics 30, 923–930. 10.1093/bioinformatics/btt656.

Lisch, D. (2013). How important are transposons for plant evolution? Nat Rev Genet 14, 49–61. 10.1038/nrg3374.

Liu, C. (2017). In Situ Hi-C Library Preparation for Plants to Study Their Three-Dimensional Chromatin Interactions on a Genome-Wide Scale. Methods Mol Biol 1629, 155–166. 10.1007/978-1-4939-7125-1_11.

Liu, S., Zhang, L., Gao, L., Chen, Z., Bie, Y., Zhao, Q., Zhang, S., Hu, X., Liu, Q., Wang, X., and Wang, Q. (2022). Differential photoregulation of the nuclear and cytoplasmic CRY1 in Arabidopsis. New Phytol 234, 1332–1346. 10.1111/nph.18007.

Love, M.I., Huber, W., and Anders, S. (2014). Moderated estimation of fold change and dispersion for RNA-seq data with DESeq2. Genome Biol 15, 550. 10.1186/s13059-014-0550-8.

Lu, C., Chen, J., Zhang, Y., Hu, Q., Su, W., and Kuang, H. (2012). Miniature inverted-repeat transposable elements (MITEs) have been accumulated through amplification bursts and play important roles in gene expression and species diversity in Oryza sativa. Mol Biol Evol 29, 1005–1017. 10.1093/molbev/msr282.

Lu, C., Kulkarni, K., Souret, F.F., MuthuValliappan, R., Tej, S.S., Poethig, R.S., Henderson, I.R., Jacobsen, S.E., Wang, W., Green, P.J., and Meyers, B.C. (2006). MicroRNAs and other small RNAs enriched in the Arabidopsis RNA-dependent RNA polymerase-2 mutant. Genome Res 16, 1276–1288. 10.1101/gr.5530106.

Lyons, M., Cardle, L., Rostoks, N., Waugh, R., and Flavell, A.J. (2008). Isolation, analysis and marker utility of novel miniature inverted repeat transposable elements from the barley genome. Mol Genet Genomics 280, 275–285. 10.1007/s00438-008-0363-0.

Matzke, M.A., Kanno, T., and Matzke, A.J. (2015). RNA-Directed DNA Methylation: The Evolution of a Complex Epigenetic Pathway in Flowering Plants. Annu Rev Plant Biol 66, 243–267. 10.1146/annurev-arplant-043014-114633.

Maumus, F., and Quesneville, H. (2014). Ancestral repeats have shaped epigenome and genome composition for millions of years in Arabidopsis thaliana. Nat Commun 5, 4104. 10.1038/ncomms5104.

Nishida, H., Ishihara, D., Ishii, M., Kaneko, T., Kawahigashi, H., Akashi, Y., Saisho, D., Tanaka, K., Handa, H., Takeda, K., and Kato, K. (2013). Phytochrome C is a key factor controlling long-day flowering in barley. Plant Physiol 163, 804–814. 10.1104/pp.113.222570.

Oki, N., Yano, K., Okumoto, Y., Tsukiyama, T., Teraishi, M., and Tanisaka, T. (2008). A genome-wide view of miniature inverted-repeat transposable elements (MITEs) in rice, Oryza sativa ssp. japonica. Genes Genet Syst 83, 321–329. 10.1266/ggs.83.321.

Pasha, A., Subramaniam, S., Cleary, A., Chen, X., Berardini, T., Farmer, A., Town, C., and Provart, N. (2020). Araport Lives: An Updated Framework for Arabidopsis Bioinformatics. Plant Cell 32, 2683–2686. 10.1105/tpc.20.00358.

Quadrana, L., Bortolini Silveira, A., Mayhew, G.F., LeBlanc, C., Martienssen, R.A., Jeddeloh, J.A., and Colot, V. (2016). The Arabidopsis thaliana mobilome and its impact at the species level. Elife 5. 10.7554/eLife.15716.

Rice, P., Longden, I., and Bleasby, A. (2000). EMBOSS: the European Molecular Biology Open Software Suite. Trends Genet 16, 276–277. 10.1016/s0168-9525(00)02024-2.

SanMiguel, P., Gaut, B.S., Tikhonov, A., Nakajima, Y., and Bennetzen, J.L. (1998). The paleontology of intergene retrotransposons of maize. Nat Genet 20, 43–45. 10.1038/1695.

Sasaki, T., Lee, T.F., Liao, W.W., Naumann, U., Liao, J.L., Eun, C., Huang, Y.Y., Fu, J.L., Chen, P.Y., Meyers, B.C., et al. (2014). Distinct and concurrent pathways of Pol II-and Pol IV-dependent siRNA biogenesis at a repetitive trans-silencer locus in Arabidopsis thaliana. Plant J 79, 127–138. 10.1111/tpj.12545.

Schmitz, R.J., He, Y., Valdes-Lopez, O., Khan, S.M., Joshi, T., Urich, M.A., Nery, J.R., Diers, B., Xu, D., Stacey, G., and Ecker, J.R. (2013). Epigenome-wide inheritance of cytosine methylation variants in a recombinant inbred population. Genome Res 23, 1663–1674. 10.1101/gr.152538.112.

Schnable, P.S., Ware, D., Fulton, R.S., Stein, J.C., Wei, F., Pasternak, S., Liang, C., Zhang, J., Fulton, L., Graves, T.A., et al. (2009). The B73 maize genome: complexity, diversity, and dynamics. Science 326, 1112–1115. 10.1126/science.1178534.

Seren, U., Grimm, D., Fitz, J., Weigel, D., Nordborg, M., Borgwardt, K., and Korte, A. (2017). AraPheno: a public database for Arabidopsis thaliana phenotypes. Nucleic Acids Res 45, D1054–D1059. 10.1093/nar/gkw986.

Sotelo-Silveira, M., Chavez Montes, R.A., Sotelo-Silveira, J.R., Marsch-Martinez, N., and de Folter, S. (2018). Entering the Next Dimension: Plant Genomes in 3D. Trends Plant Sci 23, 598–612. 10.1016/j.tplants.2018.03.014.

Stuart, T., Eichten, S.R., Cahn, J., Karpievitch, Y.V., Borevitz, J.O., and Lister, R. (2016). Population scale mapping of transposable element diversity reveals links to gene regulation and epigenomic variation. Elife 5. 10.7554/eLife.20777.

Tittel-Elmer, M., Bucher, E., Broger, L., Mathieu, O., Paszkowski, J., and Vaillant, I. (2010). Stress-induced activation of heterochromatic transcription. PLoS Genet 6, e1001175. 10.1371/journal.pgen.1001175.

Underwood, C.J., Vijverberg, K., Rigola, D., Okamoto, S., Oplaat, C., Camp, R., Radoeva, T., Schauer, S.E., Fierens, J., Jansen, K., et al. (2022). A PARTHENOGENESIS allele from apomictic dandelion can induce egg cell division without fertilization in lettuce. Nat Genet 54, 84–93. 10.1038/s41588-021-00984-y.

Wu, N., Yao, Y., Xiang, D., Du, H., Geng, Z., Yang, W., Li, X., Xie, T., Dong, F., and Xiong, L. (2022). A MITE variation-associated heat-inducible isoform of a heat-shock factor confers heat tolerance through regulation of JASMONATE ZIM-DOMAIN genes in rice. New Phytol 234, 1315–1331. 10.1111/nph.18068.

Wu, R., Lucke, M., Jang, Y.T., Zhu, W., Symeonidi, E., Wang, C., Fitz, J., Xi, W., Schwab, R., and Weigel, D. (2018). An efficient CRISPR vector toolbox for engineering large deletions in Arabidopsis thaliana. Plant Methods 14, 65. 10.1186/s13007-018-0330-7.

Xu, L., Yuan, K., Yuan, M., Meng, X., Chen, M., Wu, J., Li, J., and Qi, Y. (2020). Regulation of Rice Tillering by RNA-Directed DNA Methylation at Miniature Inverted-Repeat Transposable Elements. Mol Plant 13, 851–863. 10.1016/j.molp.2020.02.009.

Yang, G., Nagel, D.H., Feschotte, C., Hancock, C.N., and Wessler, S.R. (2009). Tuned for transposition: molecular determinants underlying the hyperactivity of a Stowaway MITE. Science 325, 1391–1394. 10.1126/science.1175688.

Zhang, H., Tao, Z., Hong, H., Chen, Z., Wu, C., Li, X., Xiao, J., and Wang, S. (2016). Transposon-derived small RNA is responsible for modified function of WRKY45 locus. Nat Plants 2, 16016. 10.1038/nplants.2016.16.

Zhang, X., and Wang, T. (2021). Plant 3D Chromatin Organization: Important Insights from Chromosome Conformation Capture Analyses of the Last 10 Years. Plant Cell Physiol 62, 1648–1661. 10.1093/pcp/pcab134.

Zhou, M., and Law, J.A. (2015). RNA Pol IV and V in gene silencing: Rebel polymerases evolving away from Pol II’s rules. Curr Opin Plant Biol 27, 154–164. 10.1016/j.pbi.2015.07.005.

