## Supplemental information for "Polymorphic Inverted Repeats near coding genes impact chromatin topology and phenotypic traits in *Arabidopsis thaliana*"

### Supplemental Figures

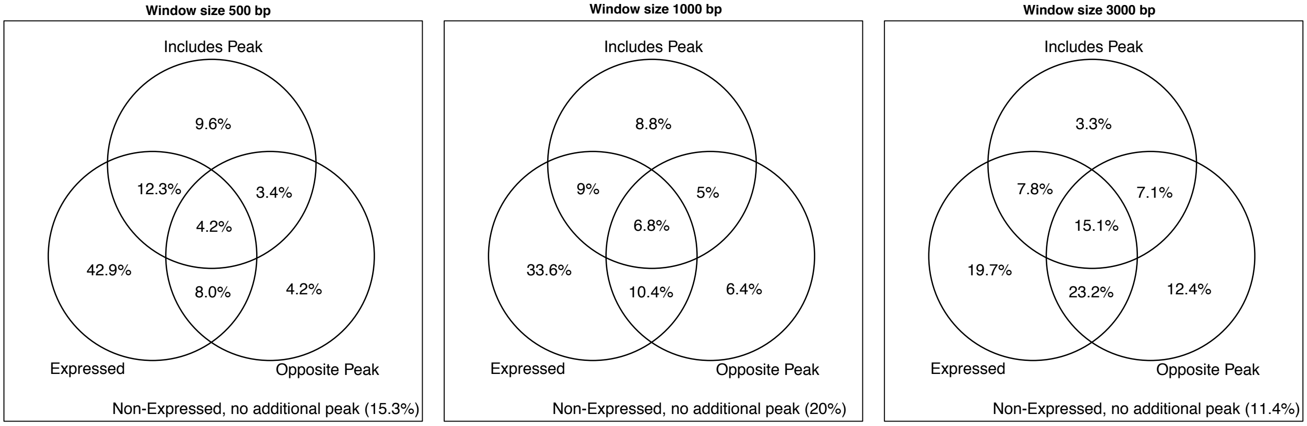


**Figure S1. IRs distribution near annotated genes in the *A. thaliana* Col-0 reference genome**

Venn Diagram showing expressed gene-hosted IRs with an additional peak of methylation nearby. Both the IR and additional methylation peaks were searched at 500, 1000 or 3000 bp windows up- and down-stream protein coding genes and displayed in separate panels. Additional methylation peaks were searched within the gene bodies (Includes Peak) or outside the opposite border of the gene relative to the IR (Opposite Peak).

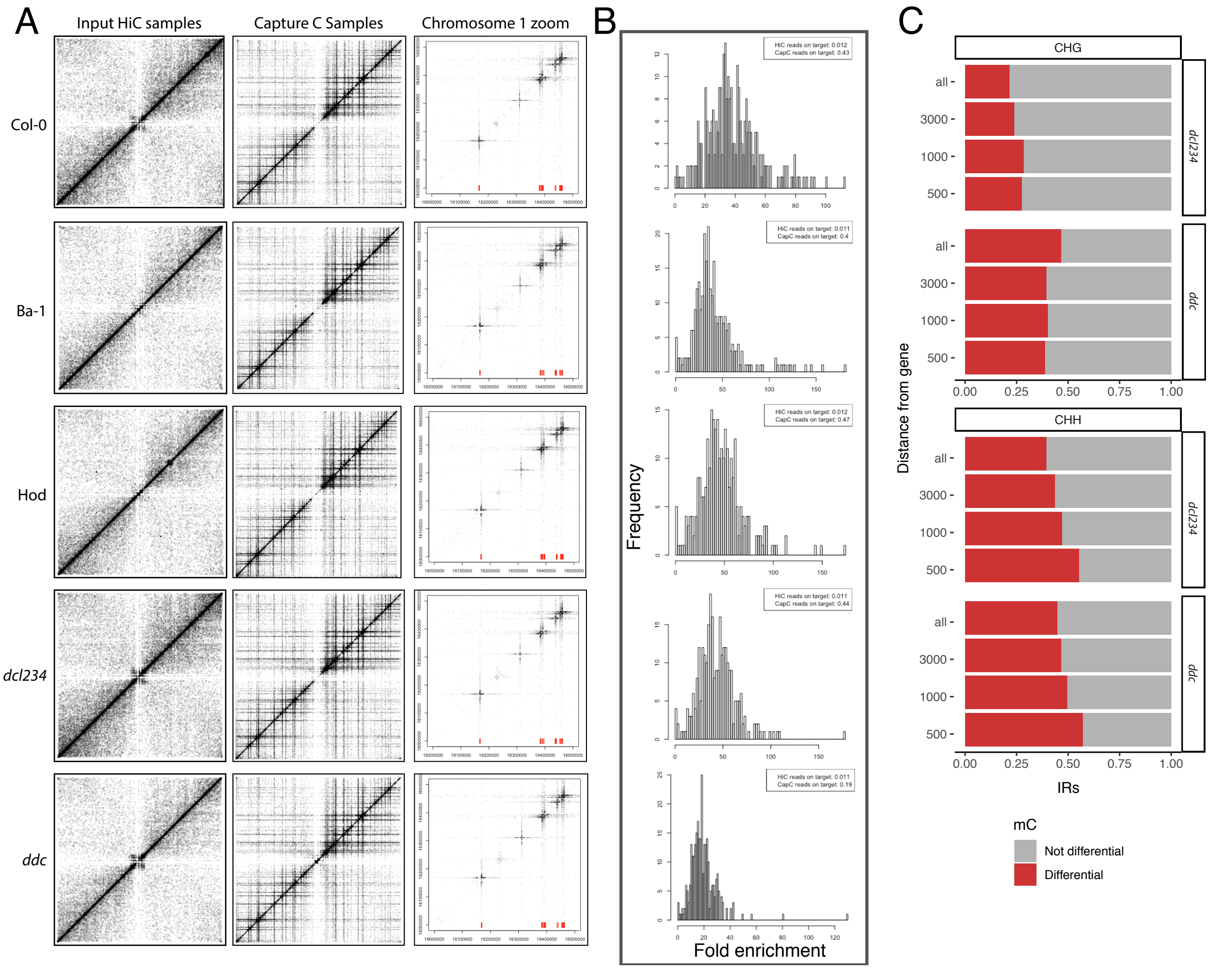


**Figure S2. Gene-hosted IRs are largely methylated**

(A) CaptureC reads distribution on chromosome 1 showing enrichment in the captured regions compared to the input HiC samples. Red segments in the right column indicate the position of six probes sets designed in a a 0.5 Mb region in Chromosome 1.

(B) Distribution of the enrichment in sequenced reads observed for each probed region in each sample relative to its original Hi-C library. The fold enrichment for each region was calculated as the ratio between the proportion of reads mapping to a region after Capture-C with respect to the proportion of reads from the sequencing of the library before capturing (HiC input). Inlay boxes on the upper right corner of each plot indicate the proportion of reads mapping to all the captured regions before and after the CaptureC enrichment.

(C) Proportion of all and gene-associated IRs showing differential DNA methylation in the CHG and CHH contexts in the *ddc* and *dcl234* mutants with respect to Col-0.

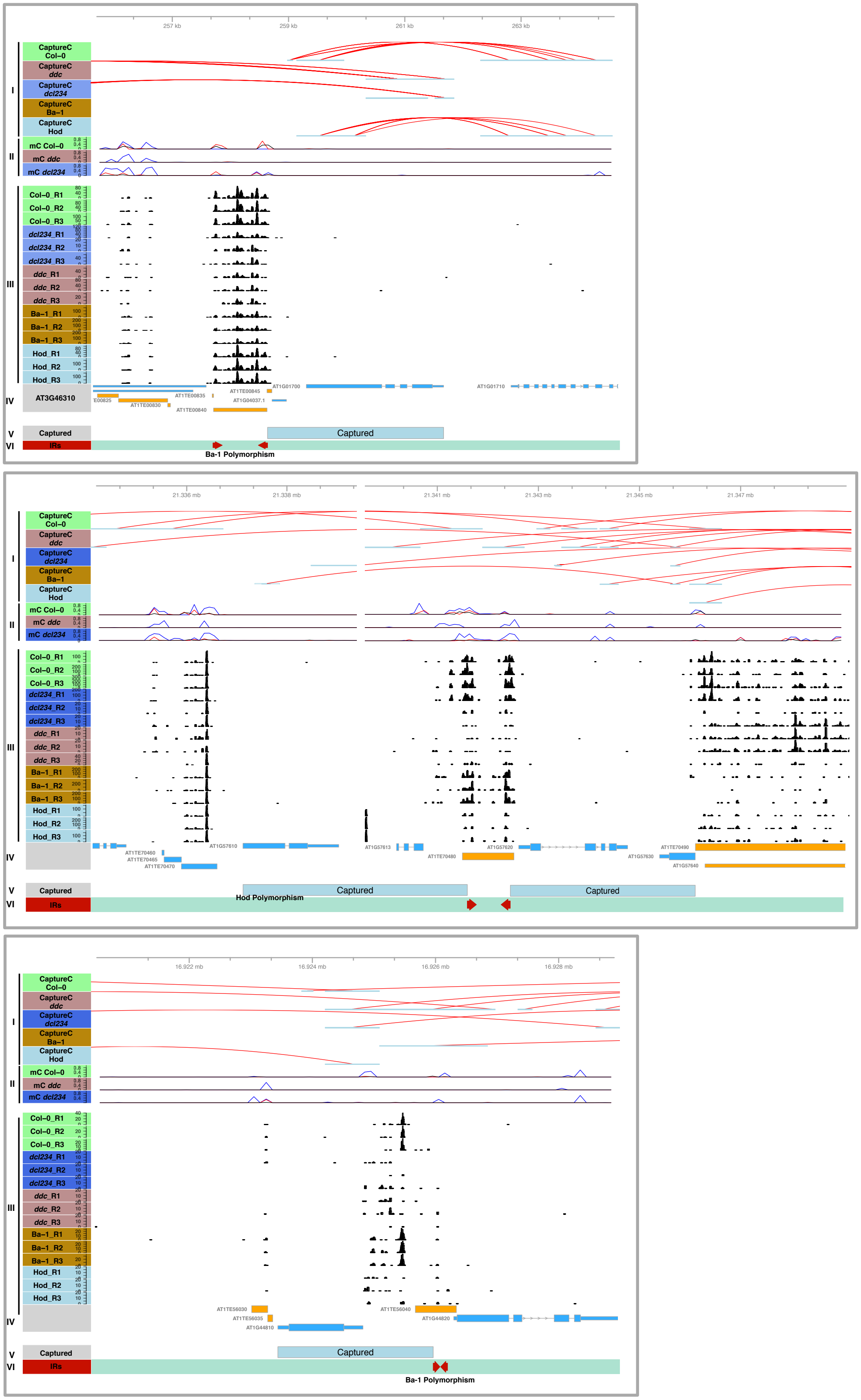

257 kb
259 kb
261 kb
263 kb
CaptureC Col–0
CaptureC ddc
CaptureC dcl234
CaptureC Ba–1
CaptureC Hod
I
mC Col–0
mC ddc
mC dcl234
II
Col–0_R1
Col–0_R2
Col–0_R3
dcl234_R1
dcl234_R2
dcl234_R3
ddc_R1
ddc_R2
ddc_R3
Ba–1_R1
Ba–1_R2
Ba–1_R3
Hod_R1
Hod_R2
Hod_R3
III
AT3G46310
IV
AT1G01700
AT1G01710
AT1TE00845
AT1TE00835
AT1G04037.1
AT1TE00830
AT1TE00840
Captured
V
IRs
VI
Ba-1 Polymorphism
21.336 mb
21.338 mb
21.341 mb
21.343 mb
21.345 mb
21.347 mb
AT1TE70460
AT1TE70465
AT1TE70470
AT1G57610
AT1G57613
AT1TE70480
AT1G57620
AT1G57630
AT1TE70490
AT1G57640
Hod Polymorphism
16.922 mb
16.924 mb
16.926 mb
16.928 mb
AT1TE56030
AT1TE56035
AT1G44810
AT1TE56040
AT1G44820

**Figure S3. Effects of IR-derived siRNAs and methylation on local genome topology**
Three selected regions of the Col-0 genome with its epigenetic and topological profiles. (I) Chromatin interactions as detected by Capture-C experiments from Col-0, Ba-1, and Hod plants or *dcl234* and *ddc* mutants. Red lines indicate interacting fragments. (II) Cytosine DNA methylation in CG (blue), CHG (red) and CHH (black) contexts. (III) 24 nt siRNAs mapping to the genomic regions as determined by sRNA sequencing in biological triplicates of wild-type Col-0, Ba-1, and Hod plants or *dcl234* and *ddc* mutants. (IV) Annotated genes in the region. (V) Region captured by the probes designed for the Capture-C experiment. (IV) Polymorphic IRs identified in this region of the genome.

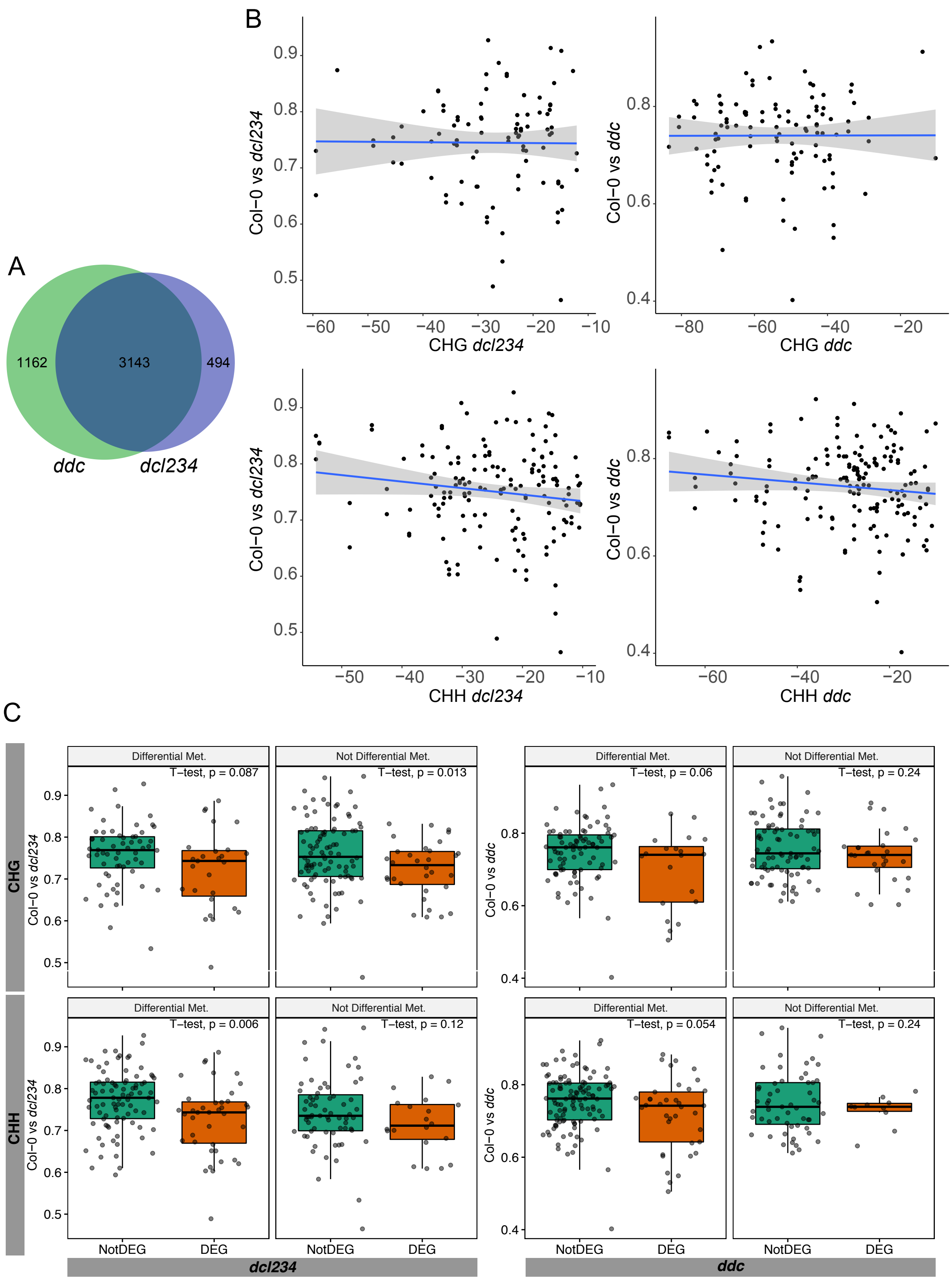


**Figure S4. Effects of IR-triggered changes on local genome topology over host genes expression**

(A) Venn diagram showing genes differentially expressed in *dcl234* and *ddc* compared to Col-0 WT plants.

(B) Scatter plots showing the structural similarity (SSIM) as calculated by CHESS linked to differentially and not differentially methylated IRs in the *ddc* and *dcl234* plants. The blue lines show the linear regression while the gray shades show the confidence interval.
(C) Box plots showing the correlation between cytosine methylation in the CHH and CHG context and structural similarity (SSIM) in differentially (DEG) and not differentially expressed (NotDEG) genes in *ddc* and *dcl234* plants.

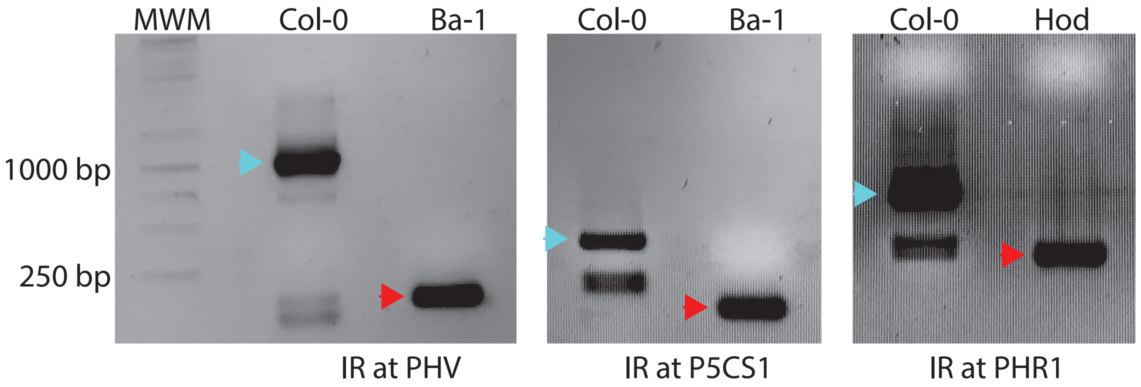


**Figure S5. Confirmation of selected IR polymorphisms in *A. thaliana* accessions**
Gel separation of amplified genomic DNA of the regions of *PHV*, *P5CS1*, and *PRH1* containing polymorphic IRs. Cyan arrowheads indicate the sequenced bands corresponding to Col-0 containing the IR sequence. Red arrowheads indicate the sequenced bands corresponding to Ba-1 or Hod accessions which lack the polymorphic IR.

**Table S1.** List of primers used in this study.

| Gene (AGI) | Sequences (5´ - 3´) | Assay |
|---|---|---|
| PHYC IR | CGATCCATCTGTTATGGGTTAAC | Genotyping |
| | AATTCAGACTTGATTAGAAAGGGG | |
| PHYC Loop | GACGTTTTGGTACATCTATCTTC | 3C-qPCR |
| | CTCATTGATGAAACTTCTTGAGG | |
| *PHYC* | CTCATTGATGAAACTTCTTGAGG | RT-qPCR |
| | GAGGGAATCGAAGAAGGCTATG | |
| PHYC (CRISPR) | GGGGAAAATCAGCCCGTAACAGTTTTAGAGCTAT GCTGAAAA | CRISP/Cas9 sgRNAs |
| | CTGTTACGGGCTGATTTTCCCCAATCACTACTTC GACTCTAG | |
| | GCTTAGTCTAGTGGTTAAACGGTTTTAGAGCTAT GCTGAAAA | |
| | CCGTTTAACCACTAGACTAAGCAATCACTACTTCG ACTCTAG | |

| | | |
|---|---|---|
| CRY1 IR | CCTTTGACAACGATTTGATATGTTG | Genotyping |
| | TTAAAGAGTATTTTCCCTTTTGTTG | |
| CRY1 Loop | CCAATAATCGTCCCACTAAAATCC | 3C-qPCR |
| | CAAGCTCGATACCAGCAGCTTG | |
| *CRY1* | GAAAGAAGCGGAGGCATAGTCCCC | RT-qPCR |
| | TGCACCTCCAAATGCACTTTTTACCC | |
| CRY1 (CRISPR) | GTCAGAATCTCTTATCTTCTTGTTTTAGAGCTATG CTGAAAA | CRISP/Cas9 sgRNAs |
| | CAAGAAGATAAGAGATTCTGACAATCACTACTTC GACTCTAG | |
| | GGTAACTATTATTCAGACACTGTTTTAGAGCTATG CTGAAAA | |
| | CAGTGTCTGAATAATAGTTACCAATCACTACTTCG ACTCTAG | |
| PHV loop1 (FW) | CACCAGAGCTTTTTCTTTACCGAG | 3C-qPCR |
| PHV loop2 (FW) | ACCGGTGAGGCCTGAGAACC | |
| PHV loop3 (FW) | GGGCAAAGACAAGCTGAGCAC | |
| PHV loop1,2,3 (RV) | TTGAATTAAAATTCAAGCTTAGCTTTG | |
| P5CS1 loop1,2 (FW) | GATGATGATGATGATGATGAAGG | 3C-qPCR |
| P5CS1 loop1 (RV) | GAGATTCTAAAGTTCAACTGACTC | |
| P5CS1 loop2 (RV) | TGTTGGACAAAGCAGTCTTATGG | |
| PHR1 loop1,2 (FW) | GCAATTGCATACATCTACTTATGG | 3C-qPCR |
| PHR1 loop1 (RV) | AGATTCTGAAAAAGCAAAGAAGGG | |
| PHR1 loop2 (RV) | TCATTTATCTGGATTTTGACTGTG | |
| Actin for EcoR1 and XbaI 3C | GTTTCTCACTTCCACATGCTATCC | 3C-qPCR |
| | CCTTGATGTCTCTCACAATTTCCC | |
| PHV | ATTTGCAAACCAAGCTGGTTTAGAC | qPCR |
| | TCGCATATCCCTGCTGCATGATC | |
| P5CS1 | AAGAGCCCCATATCAGGATTCTTCT | qPCR |
| | TGTGTAAAGACCTTCAACATCGCTC | |
| PHR1 | TGTGGCGTTCATGATCAGGGATG | qPCR |
| | GTCTCTGGTGAGCTGTTCTTCAG | |
| Actin | GGTAACATTGTGCTCAGTGGTGG | qPCR |
| | GGAGATCCACATCTGCTGGAATG | |
| PHV IR | TGTCACTATGCGCGTGAGATG | Genotyping |
| | CCAGCAAAGGCAGACCTGTC | |
| P5CS1 IR | GAAGAAGAAGAGAATGCTAGTTTC | Genotyping |
| | ACCCTAAATTCTCAAAACATTGTG | |
| PHR1 IR | ATTGATCTACGTGGATGGTTTAC | Genotyping |
| | CCATGGCTATATAGGTCAATCC | |
